# Cryosectioning-enhanced super-resolution microscopy for single-protein imaging across cells and tissues

**DOI:** 10.1101/2024.02.05.576943

**Authors:** Johannes Stein, Maria Ericsson, Michel Nofal, Lorenzo Magni, Sarah Aufmkolk, Ryan B. McMillan, Laura Breimann, Conor P. Herlihy, S. Dean Lee, Andréa Willemin, Jens Wohlmann, Laura Arguedas-Jimenez, Peng Yin, Ana Pombo, George M. Church, Chao-ting Wu

## Abstract

DNA-PAINT enables nanoscale imaging with virtually unlimited multiplexing and molecular counting. Here, we address challenges, such as variable imaging performance and target accessibility, that can limit its broader applicability. Specifically, we enhance its capacity for robust single-protein imaging and molecular counting by optimizing the integration of TIRF microscopy with physical sectioning, in particular, Tokuyasu cryosectioning. Our method, tomographic & kinetically enhanced DNA-PAINT (tkPAINT), achieves 3 nm localization precision across diverse samples, enhanced imager binding, and improved cellular integrity. tkPAINT can facilitate molecular counting with DNA-PAINT inside the nucleus, as demonstrated through its quantification of the in situ abundance of RNA Polymerase II in both HeLa cells as well as mouse tissues. Anticipating that tkPAINT could become a versatile tool for the exploration of biomolecular organization and interactions across cells and tissues, we also demonstrate its capacity to support multiplexing, multimodal targeting of proteins and nucleic acids, and 3D imaging.

## Introduction

Spatial omics technologies are advancing our understanding of the molecular principles that govern cellular function and organization^1–3^. By integrating molecular composition with spatial context, these approaches illuminate how biomolecules organize within cells and tissues. Super-resolution microscopy has expanded these capabilities, enabling visualization of biomolecules at sub-20 nm resolution^4–7^. DNA-PAINT (Points Accumulation for Imaging in Nanoscale Topography) is a single-molecule localization microscopy (SMLM) technique that achieves super-resolution imaging via transient binding of dye-labeled “imager” oligonucleotides to complementary “docking strands” attached to the target molecules^8^. DNA-PAINT enables straightforward sequential multiplexing of up to 30 targets^9–11^, single-protein resolution^12–14^, and molecular counting^15,16^, establishing it as a powerful tool for spatial biology.

The potential of DNA-PAINT relies on sample preparations that ensure accessibility to a wide range of targets while retaining cellular ultrastructure. Indeed, challenges such as fixation-induced redistribution of target molecules, antibody-induced clustering, or target loss during permeabilization can affect nanoscale imaging outcomes^17–21^. Additionally, the imaging performance of DNA-PAINT varies across sample types, molecular targets, and microscopy modalities^8,22,23^. For instance, while Total Internal Reflection Fluorescence^24^ (TIRF) microscopy offers the highest resolution for single-protein imaging with DNA-PAINT^12,14^, its axial range (∼200 nm) restricts imaging to targets near the cover glass. Most cellular targets, however, elude the accessible TIRF range and thus require alternative imaging conditions, reducing resolution^8,22,23^ and limiting its ability for counting^12,13,25–28^.

Physical sectioning offers compelling solutions to these challenges^29–32^, enabling TIRF-based SMLM imaging of cell regions otherwise inaccessible^33^ while ensuring high target accessibility and structural integrity^34,35^. Despite implementations with SMLM across diverse samples^33,36–40^, sectioning has thus far only been used for DNA-PAINT imaging of tissues^41–45^, where it is a routine step. For instance, Tokuyasu cryosectioning^46^ – known for its excellent ultrastructure preservation and antigenicity^35^ – was recently adopted for DNA-PAINT, achieving 4 nm localization precisions using TIRF and multiplexing via Exchange-PAINT^9^ on ∼350 nm rat brain cryosections without permeabilization^47,48^. Additionally, DNA-PAINT imaging of ultrathin resin sections has enabled volumetric reconstructions from sequential sections, as shown in Alzheimer’s brain tissues^45^. These studies provide compelling reasons to maximize the potential of physical sectioning for DNA-PAINT.

Here, we present “tomographic and kinetically-enhanced DNA-PAINT” (tkPAINT), a workflow that leverages physical sectioning to align sample volume with TIRF illumination, thereby greatly enhancing resolution and imager binding for robust single-protein imaging and counting. Adopting a Tokuyasu protocol for targeting RNA Polymerase II (Pol II) in HeLa cells^49^, we demonstrate the potential of physical sectioning for intranuclear DNA-PAINT imaging^22,50–54^ (**Fig. 1**a), obtaining localization precisions down to 3 nm while preserving cellular ultrastructure. We show that reducing section thickness can enhance imager binding statistics, with up to 80% of localizations attributed to Pol II signal in ∼150 nm cryosections. This enabled us to perform molecular counting with DNA-PAINT inside the nucleus. Using qPAINT^15^ (quantitative DNA-PAINT), we count antibodies within nanoscopic Pol II clusters and quantify their nuclear abundance. Extending tkPAINT to mouse tissues, we demonstrate its ability to deliver consistent conditions for single-protein imaging and counting across sample types while revealing cell- and tissue-specific heterogeneities in Pol II organization^55,56^. The versatility of tkPAINT is further highlighted through multiplexing, multimodal imaging of proteins and nucleic acids as well as 3D imaging using astigmatism. While this work pushes the capabilities of DNA-PAINT for spatial biology in single sections, we anticipate integrations of tkPAINT with well-established serial sectioning approaches^36,39,45,57^ to reconstruct larger sample volumes and entire nuclei.

**Figure 1.**
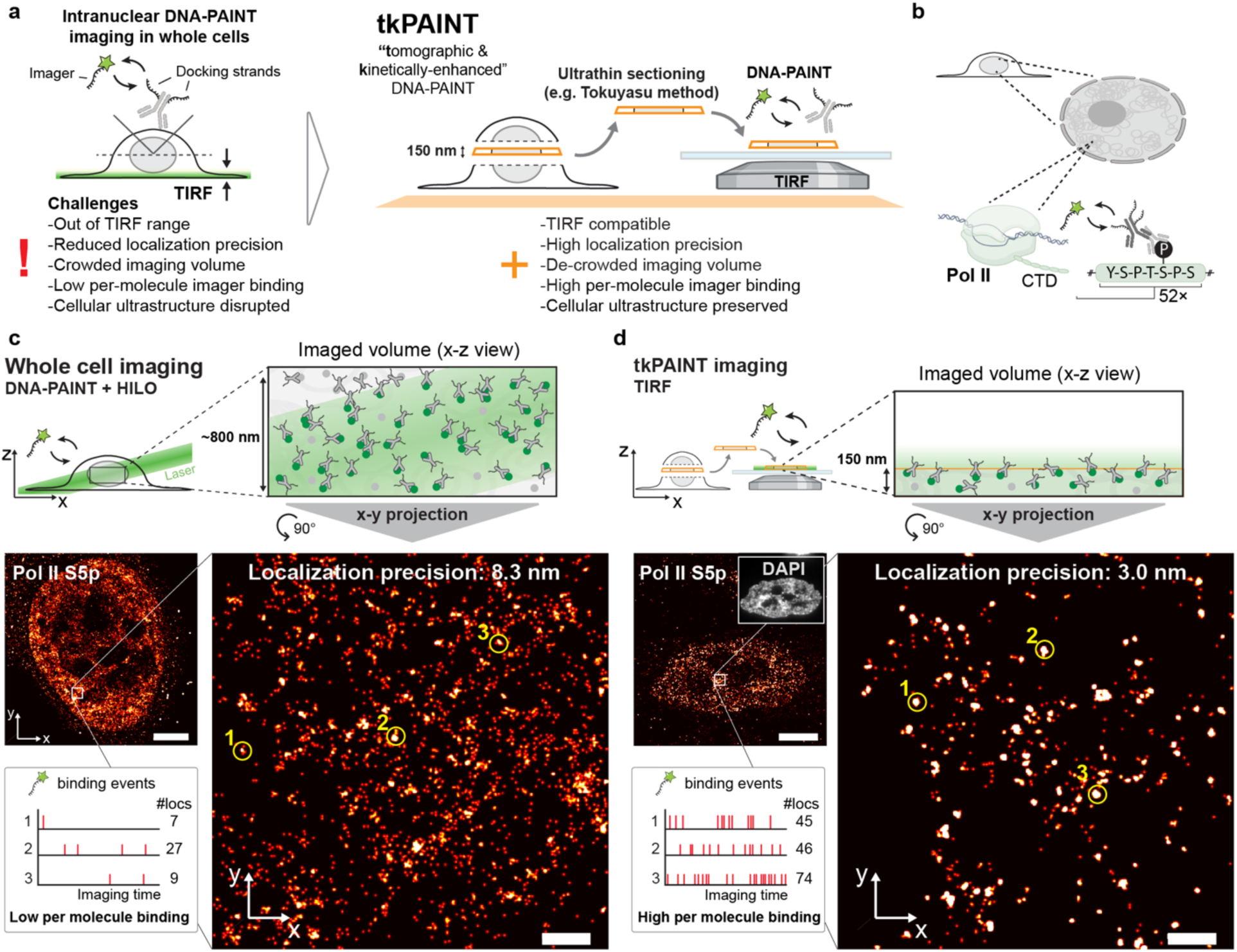
tkPAINT enables TIRF-based DNA-PAINT imaging of intranuclear targets and enhanced imager binding. **a** tkPAINT schematic. Ultrathin cryosectioning enables nuclear DNA-PAINT imaging under TIRF conditions. **b** Immunolabeling of Pol II CTD Serine-5 phosphorylation (S5p) for DNA-PAINT imaging via docking strand-conjugated secondary antibodies. **c** HILO DNA-PAINT image of Pol II S5p within whole HeLa cell. Time traces of imager binding and corresponding number of localizations are shown for the three regions indicated by yellow circles, demonstrating low per molecule binding since imager binding events are shared between a high number of labeled target molecules within the imaged volume (green circles, top schematic; grey target molecules inaccessible to antibody labeling remain unseen). **d** tkPAINT image of Pol II S5p. The inset shows the same cell imaged in the DAPI channel. Time traces of imager binding and number of localizations are shown for three regions indicated by yellow circles, demonstrating high per molecule binding. Imager binding events are shared between a low number of imaged target molecules within the imaged volume (green circles, top schematic). Scale bars, 5µm in (**c**,**d**), 400 nm in zoom-ins.

## Results

### tkPAINT enables TIRF-based DNA-PAINT throughout ultrastructure-preserved cells with enhanced resolution and binding kinetics

To develop tkPAINT, we chose to target the largest subunit of Pol II, Rpb1, a highly abundant nuclear protein. We focused, in particular, on its C-terminal domain (CTD), which features 52 heptad repeats of the consensus motif YSPTSPS, the residues of which are posttranslationally modified during transcription and are involved in promoting co-transcriptional RNA splicing^58^ (**Fig. 1**b). Using a primary antibody against hyperphosphorylated Serine-5 of the CTD (S5p), we then leveraged previously optimized protocols for diffraction-limited nuclear imaging within Tokuyasu cryosections under ultrastructure-preserving conditions^59^ (**Supplementary Fig. 1**). If not stated otherwise, we refer to ultrathin cryosections of ∼150 nm thickness as ‘cryosections’, which were used for most tkPAINT experiments presented in this work.

We labeled whole HeLa cells as well as cryosections of fixed HeLa cells with both primary antibodies and oligo-conjugated secondary antibodies designed for 2D DNA-PAINT imaging using both a classical imager^8^ (P1) and speed-optimized imager^60^ (R4) which enables faster imaging at reduced imager concentrations and hence reduced fluorescence background (**Methods**). Whole cells were then imaged using DNA-PAINT with HILO illumination, while for tkPAINT, cryosections were imaged using TIRF illumination at a TIRF angle that ensured approximately homogeneous intensity over the section thickness^61^. For HILO imaging we increased imager concentrations by 2-4-fold compared to tkPAINT, due to bleaching of diffusing imagers within the excited HILO volume, reducing the effective imager concentration. At least three datasets were acquired per condition. Duration of data acquisition was kept identical for both HILO DNA-PAINT and tkPAINT imaging and imager concentrations were adjusted individually in each experiment to ensure sparse single-molecule blinking required for obtaining localizations from individual fluorescent molecules^6^ (**Methods**).

**Figures 1**c and **1**d depict the reconstructed super-resolution images obtained via HILO DNA-PAINT and tkPAINT, respectively. Overall, localizations appeared less clustered and more widely distributed in the HILO DNA-PAINT image presumably due to the larger imaging volume crowded with antibodies and lower resolution. TIRF illumination in tkPAINT led to up to 10× higher signal-to-noise ratio, as compared to HILO (**Supplementary Fig. 2**), translating to an almost 3-fold improvement in localization precision, down to ∼3 nm as compared to ∼8.3 nm in HILO DNA-PAINT (determined via Nearest Neighbor Analysis^62^, NeNA); **Supplementary Fig. 2**). R4 enabled HILO imaging at 10x lower imager concentration compared to P1, increasing the signal-to-noise ratio by more than 4-fold. However, this did not translate to an improvement in localization precision (8.1 nm vs. 8.3 nm, respectively; **Supplementary Fig. 2**), indicating that background fluorescence from diffusing imagers had negligible influence on localization precision compared to other factors such as out-of-focus binding events, autofluorescence or optical aberrations in HILO. To confirm this, we used fluorogenic imager strands^11,63^ for HILO imaging, which suppress both fluorescence and photobleaching during diffusion, again achieving localization precisions of ∼8 nm (**Supplementary Fig. 3**).

As a reference, we performed *in vitro* DNA-PAINT imaging of surface-immobilized DNA origami^64^ structures that featured a docking strand pattern with 20 nm spacing^8^ using TIRF. This resulted in a localization precision of 2.8 nm (**Extended Data Fig. 1**), demonstrating that tkPAINT can translate the resolution achievable with TIRF under *in vitro* conditions to the nuclei of fixed cells.

Efficient nuclear antibody staining in whole cells typically requires strong permeabilization^19^, which can disrupt cellular ultrastructure, particularly in the cytoplasm^18^ (**Extended Data Fig. 2**). This disruption limits the applicability of multiplexed DNA-PAINT imaging for detergent-sensitive cytoplasmic targets, such as lysosomes^65^, alongside nuclear antigens. By enabling intracellular access through sectioning, omitting permeabilization, tkPAINT overcomes this limitation. We demonstrated simultaneous imaging of lysosome-associated membrane protein 1 (LAMP1) and RNA Polymerase II (Pol II) at sub-3 nm localization precision while preserving cellular ultrastructure, as validated by immunogold electron microscopy (**Extended Data Fig. 2**). In the nucleus, permeabilization did not lead to noticeable ultrastructural perturbation and can be used to enhance antigen accessibility throughout cryosections^66^ (**Supplementary Figs. 1 & 4**).

In addition to enhancing resolution and enabling cell-wide ultrastructural access, ultrathin sectioning inherently improves the kinetic sampling of target molecules. The reduced imaged volume allows higher per-molecule imager-binding frequency while still ensuring isolated single-molecule fluorescence events required for accurate localization (**Extended Data Fig. 3**a). In fact, inspecting individual clouds of localizations in both datasets indicated significantly higher imager binding frequencies as well as number of localizations with tkPAINT as compared to HILO DNA-PAINT (yellow circles and insets, **Fig. 1**c and d, respectively). To confirm this, we globally analyzed both datasets by dividing them into five equal temporal segments and assigning unique colors to each segment (e.g., red for the first segment, blue for the last; total imaging time ∼17 min; **Extended Data Fig. 3**b). The highly colored HILO image indicated most target molecules experienced only imager binding events during one of the time segments. In contrast, the tkPAINT image appeared predominantly white, reflecting frequent revisits of imagers to target molecules. The reduction in imaging volume with tkPAINT effectively enhances imager-binding statistics, a crucial factor in DNA-PAINT for both single-molecule profiling at high fidelity and molecular counting, as discussed in the following sections.

### tkPAINT enables nuclear imaging of Pol II at single antibody resolution and molecular counting

The repetitive binding of imagers in DNA-PAINT is a critical advantage for single-protein imaging, enabling the exclusion of localizations caused by non-repetitive imager sticking^13,67,68^. This is typically accomplished by employing clustering algorithms to identify accumulations of localizations, referred to as ‘localization clouds,’ which originate from docking strand-conjugated labels. The kinetic fingerprint of each localization cloud is then analyzed to determine whether it exhibits repetitive binding. **Fig. 2**a outlines this two-step analysis approach similar to the one by Fischer *et al.*^13^, i) applying the clustering algorithm DBSCAN^69^ to detect localization clouds and ii) using a kinetic filter to exclude clouds that lack repetitive binding and are likely attributable to non-specific imager sticking (for detailed analysis steps and parameters see **Supplementary Fig. 5**).

**Figure 2.**
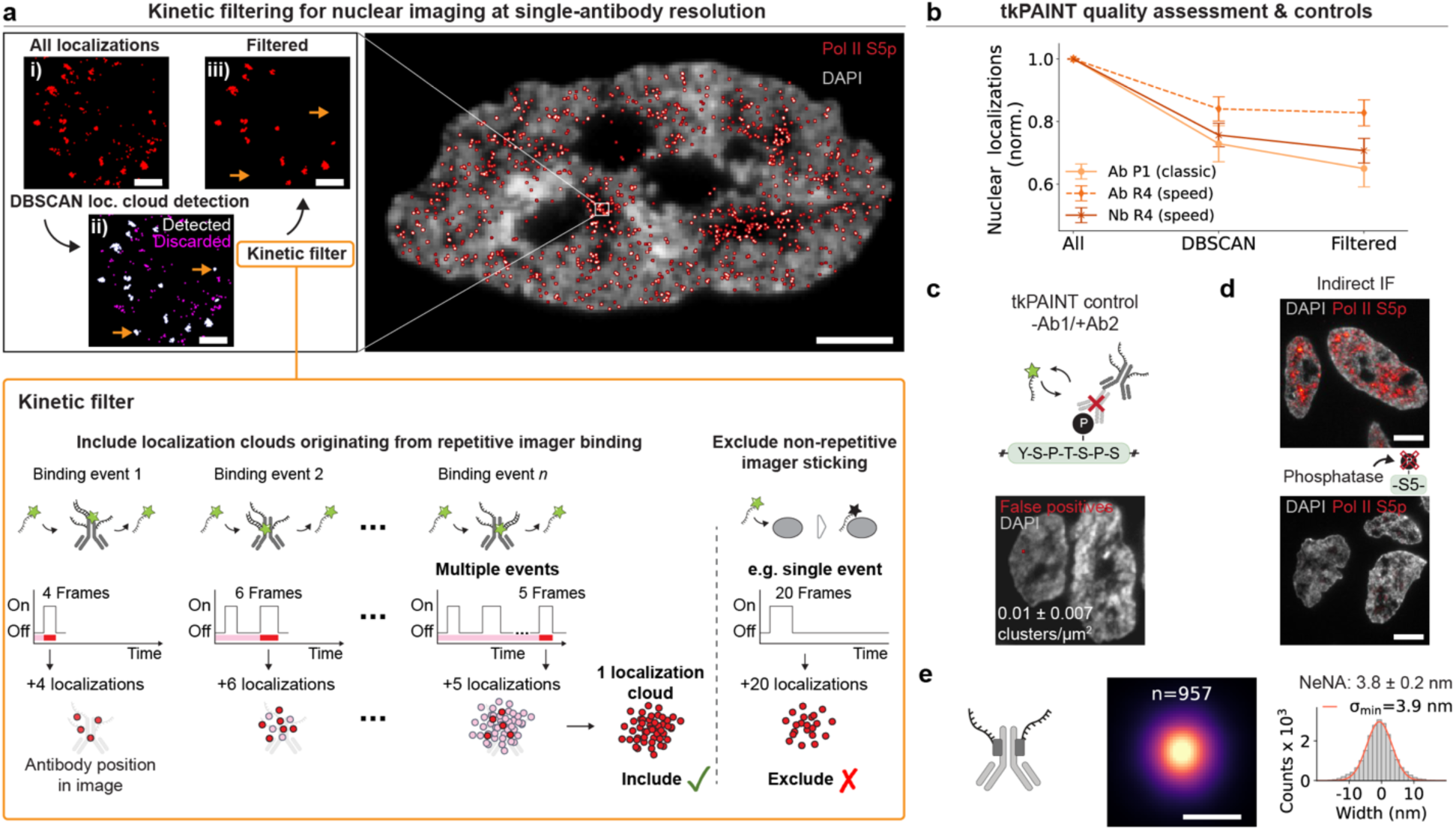
Nuclear imaging of Pol II at single antibody resolution via tkPAINT. **a** Schematic of tkPAINT data processing. i) image showing all raw localizations. ii) DBSCAN clustering is applied to detect localization clouds (white) and remaining localizations are discarded (magenta). iii) A kinetic filter^67,68^ is applied to discard localization clouds originating from non-repetitive imager sticking (orange arrows in ii) and iii); schematic description in orange box below and detailed in Supplementary Fig. 5). **b** Kinetic filter yield shown for tkPAINT Pol II S5p datasets imaged using different imager sequences (P1 – classic^8^ vs. R4 – speed^60^) or different secondary labeling strategies: secondary antibody (Ab) vs. secondary nanobody (Nb). Normalized localization counts with respect to all initial nuclear localizations showing relative loss of localizations in each analysis step. **c** tkPAINT negative control imaging of sample processed with the standard staining protocol but leaving out primary antibody and incubating secondary antibodies only. Mean and standard deviation of number of false positive localization clusters per nuclear area displayed. **d** Top: Diffraction-limited indirect immunofluorescence image of cryosections labeled for Pol II S5p (red) and DAPI (white). Bottom: same as left, but cryosections were treated with phosphatase prior to immunostaining. **e** Resolution benchmarking for Pol II S5p datasets based on secondary nanobody labeling. Middle: rendered sum images of center-of-mass aligned smallest identifiable localization clouds (see Supplementary Fig. 8 for details and additional datasets. Number of localization clouds in sum image stated above). Right: histogram showing the corresponding distribution of localizations fitted with a Gaussian (red curve) to obtain its standard deviation *σ*_min_. Scale bars, 5 µm in (**a,e**), 3 µm in (**d**) and 10 nm in sum image in (**e**).

For the tkPAINT datasets imaged with speed-optimized imager R4, over 80 % of nuclear localizations were identified as repetitive localization clouds, demonstrating efficient and targeted imaging (**Fig. 2**b). In comparison, classic imager P1 yielded a post-filtering rate of 60 %, consistent with the expected benefits of speed-optimized imagers^60,70^. Since multiple secondary antibodies can bind to a single primary antibody, this may amplify imager binding at individual target epitopes. To assess this, we repeated tkPAINT imaging using R4-conjugated secondary nanobodies, which limit the number of docking strands to a maximum of two per primary antibody. We indeed observed a minor reduction compared to R4-labeled secondary antibodies, however, still providing an excellent post-filtering localization yield of ∼70 %.

However, repeated imager binding on its own is not necessarily indicative of specificity since intrinsic cellular features could potentially also lead to repeated binding. Furthermore, secondary labels could non-specifically bind and thus position docking strands within the sample. To estimate the impact of false positive localization clouds, we performed a set of negative controls under conditions identical to that of previous tkPAINT acquisitions, but on cryosections that were incubated only with secondary antibody/nanobody and no primary antibody (**Fig. 2**c). For all tkPAINT imaging conditions, we found a negligible contribution (∼1 %) of false positive localization clouds in both cases as compared to tkPAINT experiments labeled with both primary and secondary antibody/nanobody (**Fig. 2**c and **Supplementary Fig. 6**). Increasing section thickness led to higher localization cloud densities, however, as expected both resolution and kinetic enhancement diminished (**Supplementary Fig. 7**). Thinner cryosections (∼80 nm) as used in our immunogold electron microscopy experiments (**Extended Data Fig. 2**) yielded sparse antibody signal and required delicate handling, making 150 nm our default thickness for tkPAINT. Lastly, we tested the specificity of the primary antibody against Pol II S5p by treating cryosections with phosphatase in order to neutralize phosphorylation sites prior to staining for indirect immunofluorescence^59^. Reassuringly, this led to a 3-fold signal loss (**Fig. 2**d).

The sparse distribution of localization clouds after kinetic filtering in tkPAINT (**Fig. 2**a) was reminiscent of immunogold experiments in which antibodies labeled with gold nanoparticles (diameters ∼5-15 nm) permit antigens to be detected in cryosections by TEM at the level of single antibodies^35^. To determine whether the resolution possible through tkPAINT would enable single antibodies to be visualized, we performed a range of center-of-mass alignments to obtain averaged sum images for a decreasing minimum number of localizations per cloud (**Supplementary Fig. 8**). We found that localizations in sum images were approximately Gaussian distributed with their standard deviations converging to a minimum. In other words, further reduction of localizations per cloud did not reduce the localization spread. **Figure 2**e displays a convergent sum image with a standard deviation (*σ*_min_) of ∼3.9 nm and a full width at half maximum of ∼9 nm (FWHM ≈ 2.355×*σ*_min_), indicating that localizations likely accumulated from individual antibodies, whose physical size is ∼10 nm^71^. Since secondary-nanobody labeling reduces the total label size, localization clouds more accurately reflect underlying Pol II S5p epitope positions compared to the increased label size of secondary antibody labeling (**Supplementary Fig. 6**). Together with the controllable number of docking strands per primary antibody (up to two) we thus focused our efforts with tkPAINT to quantify nuclear Pol II S5p based on secondary nanobody labeling (R4).

Since several antibodies can likely bind to a single CTD or to several Pol II molecules close by, we asked whether we could exploit the enhanced imager binding kinetics in tkPAINT to count the number of Pol II antibodies in larger localization clouds (**Fig. 3**a). In qPAINT^15^ (quantitative DNA-PAINT), the average imager binding frequency for the smallest identifiable localization clouds in a dataset is taken as a reference (**Fig. 3**b). Assuming the reference represents single antibodies, a localization cloud with *N* antibodies would have an *N*-times higher binding frequency^15^.

**Figure 3.**
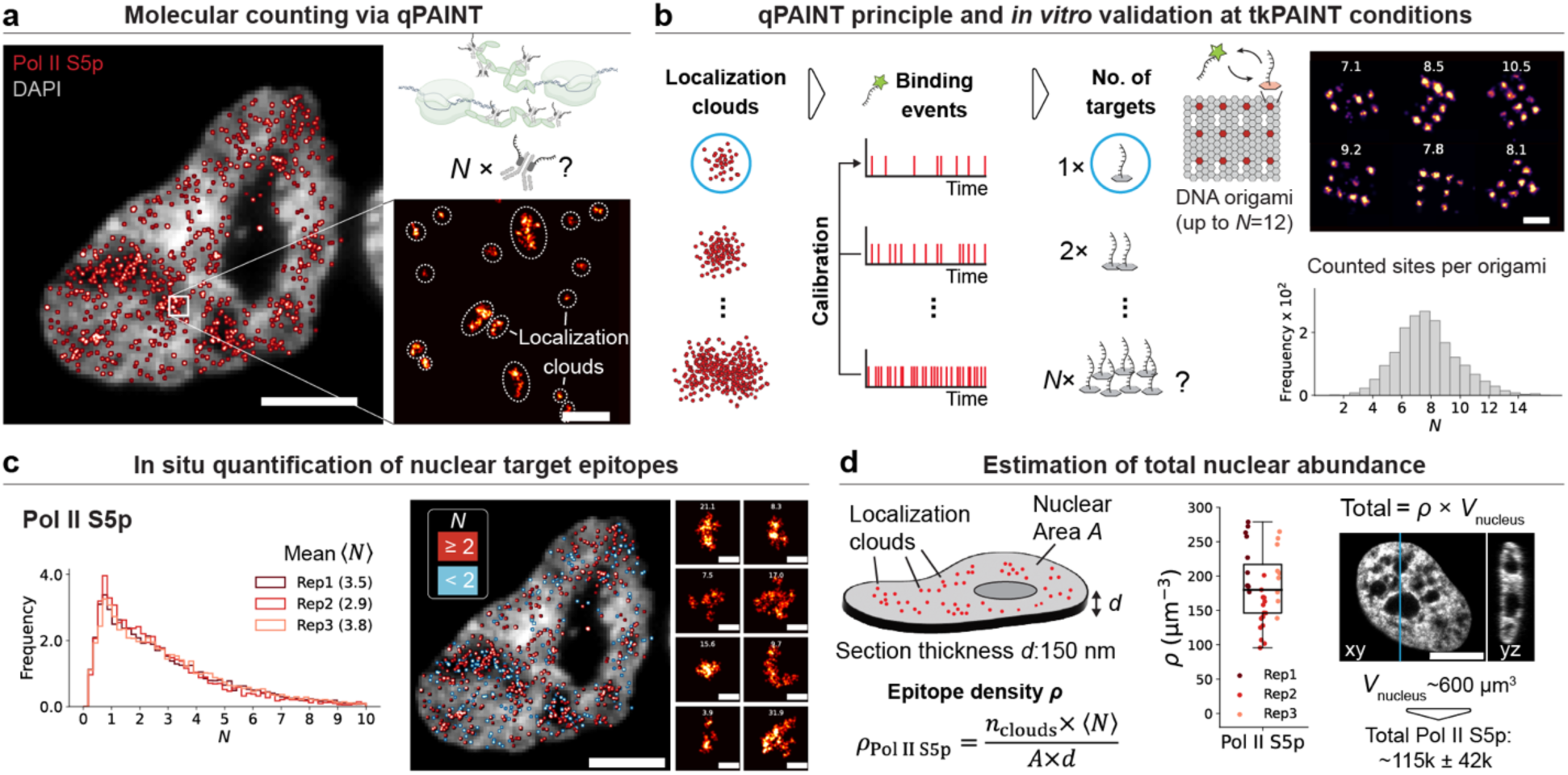
Counting Pol II antibodies in nanoscopic complexes and total nuclear abundance. **a** tkPAINT Pol II S5p datasets contain heterogeneous localization clouds and qPAINT can be exploited to count antibodies localization cloud. **b** Validating the qPAINT using DNA origami imaged under tkPAINT conditions. Left: qPAINT principle^15^. The smallest identifiable localization clouds, i.e. single docking strands on DNA origami (blue circle), serve as a calibration to measure the average imager binding frequency. Relative counting can be performed by comparing the binding frequency of each individual localization cloud to the calibration binding frequency. Right: Distribution of docking strand counts per DNA origami obtained via qPAINT analysis. Right: including six exemplary origami with their respective counting result stated above. **c** Left: Distribution of antibody counts per localization cloud for three Pol II S5p tkPAINT datasets obtained via qPAINT analysis. Right: image displayed in (a), re-rendered by coloring localization clouds according to their antibody counts: red (≥2), blue (<2). The small images on the right shows six exceptionally large localization clouds with antibody counts stated above. **d** Physical sectioning enables straightforward quantification of total nuclear target abundance by calculating the antibody density for the target epitope *ρ*_Pol II S5p_ by knowing the number localization clouds, the average number of antibodies per localization cloud. the nuclear area and the section thickness. *ρ*_Pol II S5p_ for the three datasets in (c) is shown. Measuring the average nuclear volume of intact HeLa cells via confocal microscopy enables estimation of total nuclear abundance (x-y and x-z representation shown along the slice indicated by the blue line. The volume was averaged over 15 nuclei). Scale bars, 3 µm in (**a**,**c**,**d**), 100 nm in zoom-in in (**a**) and 40 nm in zoom-ins in (**b**,**d**).

We first turned to DNA origami featuring up to 12 DSs to validate the applicability of qPAINT analysis under our experimental tkPAINT conditions. Furthermore, on DNA origami single DSs can be unambiguously chosen as reference clouds. **Fig. 3**b displays the counting results obtained via qPAINT analysis, confirming the expected number of on average ∼8 DSs per origami which was in good agreement with visual inspection (see **Supplementary Fig. 9** for additional 400 randomly selected origami and analysis schematic), confirming our ability to perform molecular counting.

In tkPAINT Pol II S5p datasets most sparse localization clouds likely corresponded to single antibodies according to the previously observed spatial localization spread. We thus performed qPAINT analysis using Pol II S5p localization clouds with a convex hull area smaller than the 20th percentile as the qPAINT reference (see **Supplementary Fig. 10** for a detailed analysis schematic). Comparing the imager binding frequency of single docking strands on DNA origami with the one measured for single antibodies in tkPAINT datasets we obtained on average ∼1.6 bound nanobodies per primary antibody (**Supplementary Fig. 10**).

**Figure 3**c displays the counting results (*N*) obtained from three independent experiments, each with a prominent single antibody peak and a decreasing tail of localization clouds containing higher numbers of antibodies. The *N* distributions were in close agreement, with on average 3.2 ± 0.4 antibodies per localization cloud. Localization clouds containing tens of antibodies indicated hot spots of active Pol II (**Fig. 3**c). Based on these counting results and the known cryosection dimensions, we measured an average nuclear antibody density of 165 ± 45 µm^-3^ (**Fig. 3**d). This translated to ∼115,000 ± 42,000 Pol II S5p antibodies per nucleus, which aligns with earlier estimates of ∼65,000 engaged Pol II^72^ and ∼320,000 copies of Rpb1^73^ per HeLa cell.

It is likely that our quantification underestimated the true abundance of phosphorylated S5 (in theory up to 52x per CTD) due to steric effects. Smaller primary labels such as nanobodies against S5p could further improve quantifications and reduce linkage errors. Notably, we could not determine whether the CTD of one or multiple Pol II molecules is present in a localization cloud; however, future studies using C-terminally tagged Pol II cell lines^74^ could allow to address this question.

Finally, our data enabled us to assess the spatial distribution of Pol II S5p, which is known to associate with active chromatin or nuclear compartments, such as transcription factories and nuclear speckles^58^. Such nuclear regions correlate with low intensities when stained for DNA using DAPI (4ʹ,6-diamidino-2-phenylindole). Indeed, we observed both higher antibody counts (**Fig. 3**c) and a higher overall localization cloud density as determined by nearest neighbor distance analysis^75^ for these regions (**Extended Data Fig. 4**).

### Resolution and kinetic enhancement translate to tkPAINT imaging in mouse tissues

Encouraged by successful applications of DNA-PAINT to semi-thin (∼350 nm) Tokuyasu tissue sections^47,48^, we hypothesized that tkPAINT could also enable kinetic filtering, single-antibody resolution and molecular counting in tissue samples. To test this, we prepared two mouse tissue types (cerebellum and spleen) following established protocols^76^ (**Methods**) and processed 150 nm cryosections for tkPAINT tissue imaging of Pol II S5p, using R4-secondary nanobodies. Figure 4a depicts super-resolved tkPAINT images of Pol II S5p within cerebellum and spleen cryosections (two datasets were acquired per tissue type). As expected^47^, we obtained similar localization precisions as previously in HeLa sections (∼3 nm). The kinetic enhancement enabled by physical sectioning also translated to tissue imaging: ∼55 % of nuclear localizations could be assigned to repetitive localization clouds, confirming the suitability of tissue data for quantitative analysis (**Fig. 4**b). The lower kinetic filtering yield compared to HeLa (∼70 %) indicated slightly elevated sticking of R4 in both tissue types. We also performed center-of-mass alignments to obtain averaged sum images with decreasing minimum number of localizations per cloud to find the converging distribution width (**Supplementary Fig. 11**), further confirming tkPAINT’s capability for single-antibody resolution in tissues (*σ*_min_≈3.4 nm and FWHM≈8 nm; **Fig. 4**c).

**Figure 4.**
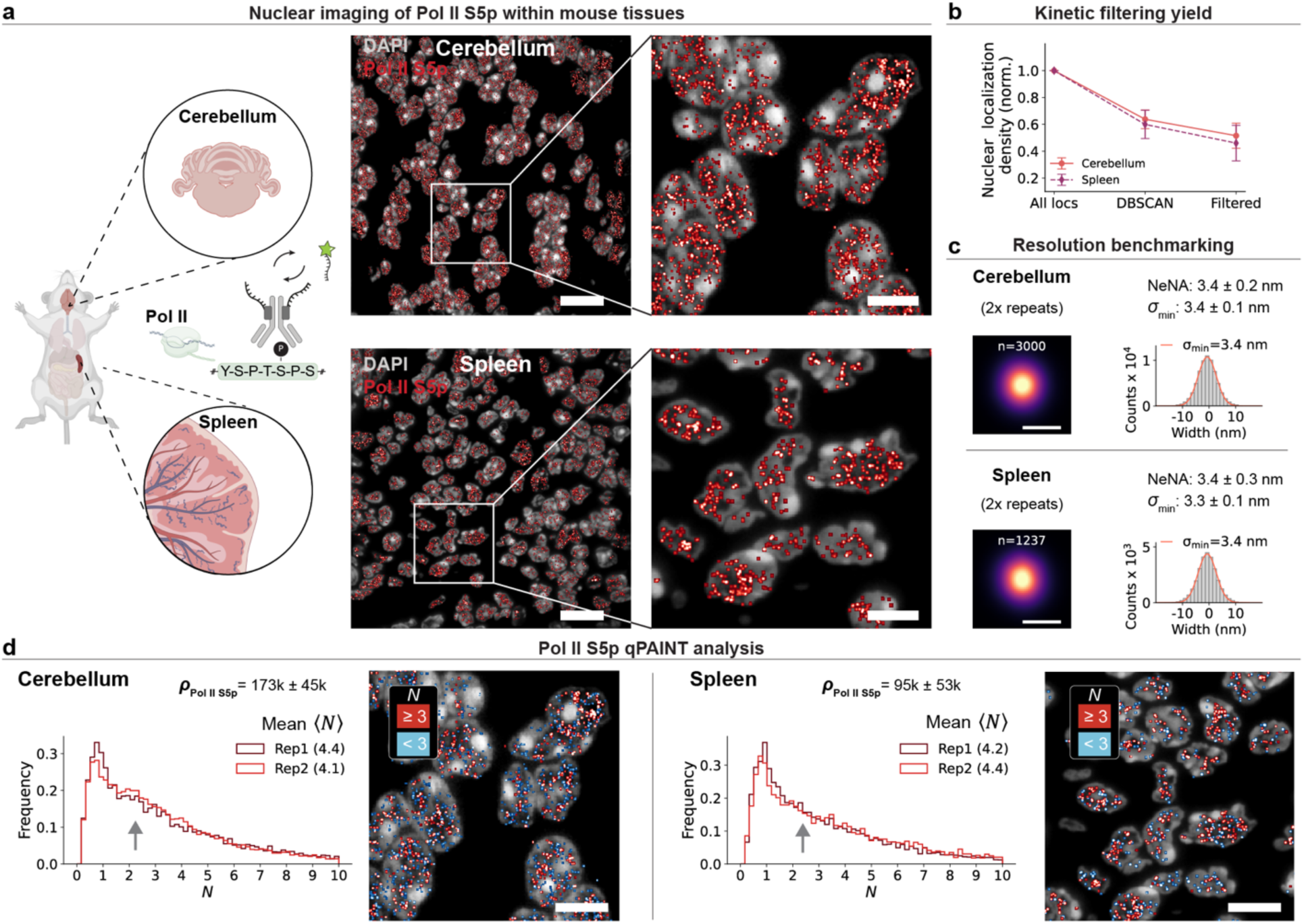
Single-protein resolution and kinetic enhancement translate to tkPAINT tissue imaging. **a** Tissue blocks of mouse cerebellum and spleen were processed for Tokuyasu sectioning and subsequently stained for Pol II S5p prior to imaging. Top: tkPAINT image of region of mouse cerebellum and zoom-in to white box. Bottom: tkPAINT image of a region of mouse spleen and zoom-in to white box. **b** Kinetic filtering yield for cerebellum and spleen tkPAINT datasets (2x each). Normalized localization counts with respect to all nuclear localizations showing relative localizations loss in each analysis step. **c** Resolution benchmarking for cerebellum and spleen tkPAINT datasets (top and bottom, respectively). Left: rendered sum images of center-of-mass aligned localization clouds for one data set each. Right: histograms showing the corresponding distribution of localizations fitted with a Gaussian (red curve) to obtain its standard deviation *σ*_min_ (see Supplementary Fig. 7 for details and Supplementary Fig. 10 for all datasets). **d** qPAINT distributions showing number of Pol II S5p antibodies per localization cloud for cerebellum and spleen datasets. The antibody density *ρ*_Pol II S5p_ is stated above. Scale bars, 10 µm in (**a**), 5 µm in zoom-ins in (**a**) and 10 nm in (**c**).

Finally, we performed qPAINT analysis to obtain spatially-resolved antibody counts in the nuclei of both tissue types (**Fig. 4**d). Each tissue yielded reproducible qPAINT distributions and averaged ∼4.2 antibodies per localization cloud, higher than the ∼3.2 observed in HeLa cells for the Pol II S5p epitope (**Fig. 3**c). Interestingly, while the qPAINT distribution for spleen closely matched that of HeLa cells, with a single antibody peak and a long tail of higher antibody counts, the cerebellum datasets showed a second peak at ∼2.5 antibodies (arrows, **Fig. 4**d). We observed a nearly two-fold enriched average nuclear density in cerebellum nuclei (95 ± 53 µm^-3^ and 173 ± 45 µm^-3^) and a higher cell-to-cell variability in spleen cells. These findings might reflect an intrinsic heterogeneity of transcriptional activity between tissue types and/or a higher number of cell types within the spleen (**Fig. 4**d). Furthermore, our results demonstrate that tkPAINT provides consistent imaging performance across diverse sample types and paves the way for probing molecular organizations between cultured cells and tissues.

### Multiplexed and multimodal tkPAINT for nuclear nanoscale imaging in 2D and 3D

Next, we turned our attention toward several proof-of-concept demonstrations, showcasing the versatility of nuclear tkPAINT imaging with respect to multiplexed single-protein imaging. Circumventing use of any secondary label for Exchange-PAINT^47,77^, we conjugated primary antibodies targeting the nuclear lamina (Lamin A/C) and nuclear speckles (SC35^78^), each with an orthogonal docking strand sequence in order to enable multiplexed imaging by sequential exchange of the complementary imager strands for each imaging round (**Fig. 5**a). Exchange-PAINT has the advantage of being free of chromatic aberrations since all imaging rounds can be acquired in the same color channel^9^. Figure 5b shows a multiplexed Exchange-tkPAINT image of Lamin A/C, Pol II S5p and SC35, sequentially imaged and subsequently reconstructed using pseudo colors. Not only did sequential imaging enable us to perform quantitative analysis for all three nuclear antigens in parallel, it permitted the spatial probing of intermolecular relationships and features. **Extended Data Fig. 5** provides an overview on how multiplexed tkPAINT data can aid the study of nuclear organization. For example, we observed two peaks in the distribution of nearest neighbor distances for Lamin A/C, which allowed us to separate the signal into a nucleoplasmic and lamina-association fraction^79,80^. Measuring nearest neighbor distances between Pol II S5p and SC35 indicated a spatial organization of Pol II S5p around nuclear speckles with SC35 at their center, as previously observed with TSA-Seq^81^.

**Figure 5.**
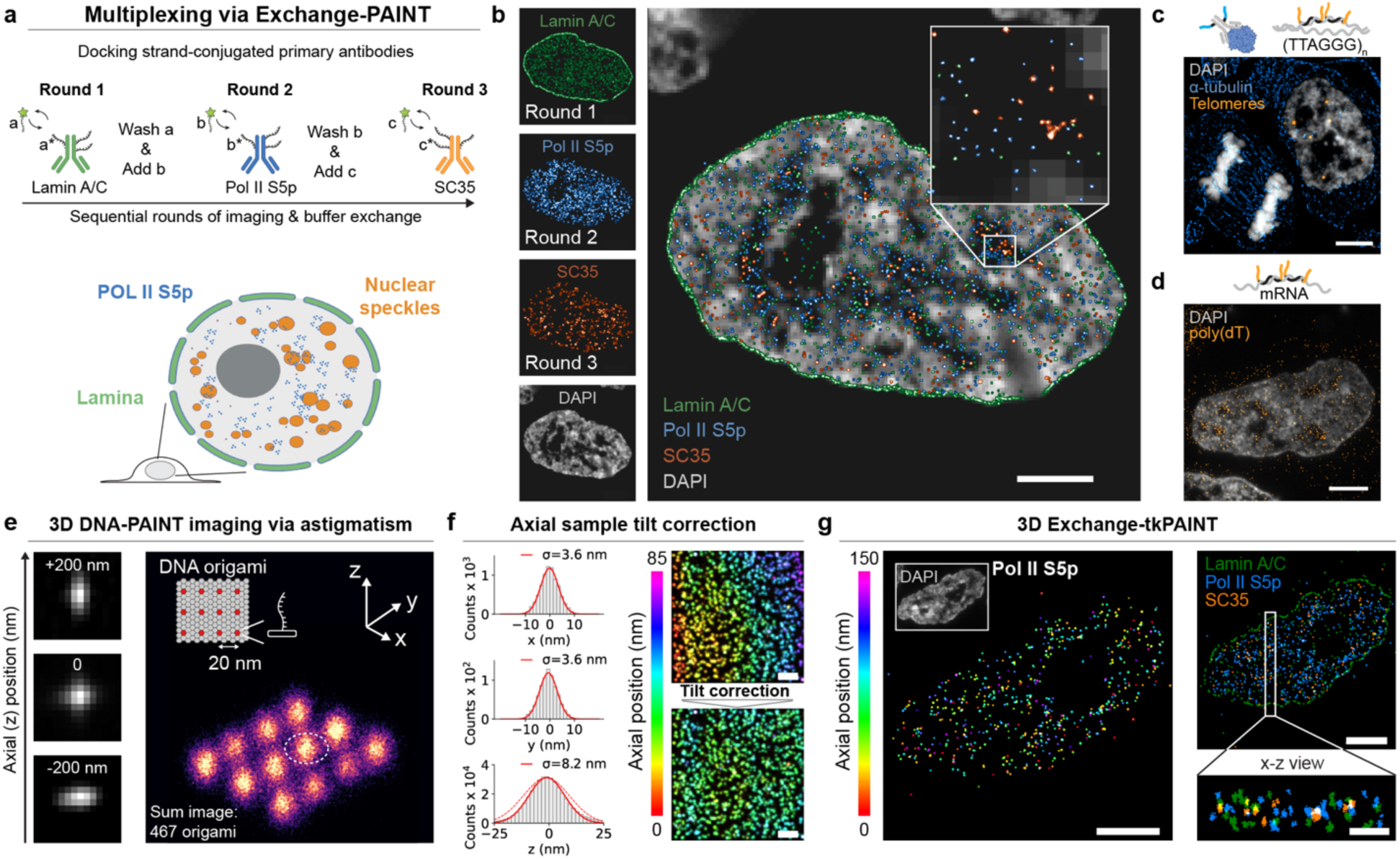
Multiplexed and multimodal nuclear nanoscale imaging in 2D and 3D. **a** Schematic of Exchange-tkPAINT targeting Lamin A/C, Pol II S5p and SC35. Primary antibodies are conjugated with orthogonal docking strands (sequences *a\**, *b\** and *c\**) and sequentially imaged with imager strands *a*, *b* and *c*. Previous imager is washed out and subsequent imager is added between rounds of imaging. **b** Multiplexed tkPAINT image reconstructed from three rounds of sequential imaging. **c** Combined imaging of protein and DNA using tkPAINT targeting of α-tubulin and telomere repeats via FISH. **d** tkPAINT imaging of mRNA via poly(dT) hybridization probes. **e** Validation of cylindrical lens addition to a commercial TIRF system for 3D DNA-PAINT imaging. Left: Astigmatism-based encoding of axial position by reshaping the point spread function. Right: Sum DNA-PAINT 467 DNA origami with 20 nm docking strand pattern. **f** Left, top and middle: x and y line plot histograms across docking strand position indicated by white dashed circle in (a). Left, bottom: Axial distribution of z coordinates of DNA origami data set. The standard deviation obtained by a Gaussian fit (red curve) is given above the histograms. Right: correction of axial sample tilt affecting measured z-distributions (red dashed curve in c). **g** Left: 3D tkPAINT of Pol II S5p. The color code indicates axial position of antibody signal over a range of 150 nm. Right: 3D Exchange-tkPAINT image of Lamin A/C (green), Pol II S5p (blue) and SC35 (orange). Side view (x-z) of localization clusters projected from white box. Scale bars, 5 µm in (**b**), 3 µm in (**c**) and 150 nm in zoom-in.

Beyond multiplexed protein imaging, a potentially even more powerful aspect of cryosections is that the same sections can be subject to both immunostaining and fluorescence in situ hybridization^82,83^, enabling analyses of the interplay between targeted proteins and specific sequences of RNA and/or DNA. Here, we performed proof-of-principle tkPAINT imaging of α-tubulin in cryosections that had additionally been labeled for telomeric repeats via in situ hybridization (**Fig. 5**c). Similarly, hybridization of a poly(dT) probes enabled us to perform tkPAINT imaging of mature mRNA (**Fig. 5**d).

TkPAINT data, previously generated through 2D imaging and, thus, resembling a two-dimensional projection of molecules within cryosections, could be significantly enhanced by accessing the axial dimension for a true interrogation of nanoscale organization. To this end, we constructed a simple and affordable (∼700$) custom addition to our commercial TIRF system that allowed us to insert a cylindrical lens in front of the camera for astigmatic 3D imaging^84^ (**Supplementary Fig. 12**). We first benchmarked our 3D imaging capability, again using surface-immobilized DNA origami with 20-nm docking strand spacing. Although the docking strand arrangement, itself, was in 2D, it nevertheless allowed us to determine the achievable axial resolution in z as well as assess whether astigmatism would significantly reduce our lateral resolution. Figure 5e shows an averaged 3D DNA-PAINT sum image of (∼450 origami), demonstrating that individual docking strands could be laterally visualized at FWHM_x,y_≈8.5 nm (σ_x,y_ ≈3.6 nm), which was sufficient to resolve the 20-nm-spaced pattern (**Fig. 5**f). During DNA origami experiments, we observed that glass slides could be tilted with respect to the optical axis, as revealed when we colored localizations according to their axial position (**Fig. 5**f). To account for this tilt, we performed a z-correction by fitting and subtracting a 2D plane^85^ (**Fig. 5**f and **Supplementary Fig. 13**). Post tilt-correction, 3D DNA-PAINT imaging of DNA origami yielded an axial distribution of localizations at FWHM_z_≈20 nm (σ_z_ ≈8.5 nm), in line with the known ∼2× axial resolution drop for astigmatic 3D SMLM^84^. An axial resolution of 20 nm would nevertheless allow us to determine distinct axial positions of antibodies within cryosections with a thickness of ∼150 nm.

These validations enabled us to move on to 3D tkPAINT imaging within cryosections of fixed HeLa cells, repeating sequential imaging of Lamin A/C, Pol II S5p, and SC35 (**Fig. 5**g). The left image in Figure 5g shows the super-resolved Pol II S5p image rendered with a range of colors according to the z-position of each localization over an axial range of 150 nm. It has been shown that, for unpermeabilized cryosections, antibody labeling happens predominantly at both surfaces of sections^86^. However, the permeabilization step in our protocol ensured antibody penetration throughout the sections, as seen for both localization clouds of all colors in the Pol II S5p image alone and the x-z projection of the multicolor Exchange-tkPAINT image (**Fig. 5**g, left and right, respectively; see **also Supplementary Fig. 4**). Overall, our 3D tkPAINT results are in close agreement with the cryotome setting for a cutting thickness of 150 nm. Measuring the overall z-distributions, we observed that while Lamin A/C and Pol II S5p labeling penetrated more homogeneously, SC35 exhibited stronger staining toward the top half of the section (**Supplementary Fig. 14**). This result reinforces the additional benefit of using smaller labels such as primary nanobodies in the future.

## Discussion

With tkPAINT, we used ultrathin sectioning to align sample volume with TIRF illumination, maximizing the capability of DNA-PAINT for single-protein imaging and counting across diverse samples and molecular targets. By leveraging the Tokuyasu method^46^, we overcame the range constraint of TIRF to access distal intracellular regions^33^, such as the nucleus, and demonstrated tkPAINT imaging throughout ultrastructure-preserved HeLa cells down to 3 nm localization precision. For imaging nuclear antigens such as Pol II, this enabled up to three-fold improved resolution as compared to HILO imaging in whole cells. Physical sectioning not only enhanced resolution and antigen accessibility but also de-crowded the sample volume, improving imager binding statistics critical for robust single-protein imaging^13,14^ and counting^15,68^. This allowed us to count antibodies within nanoscopic Pol II clusters as well as to quantify cell- and tissue-specific heterogeneities in Pol II organization. Additionally, sequential multiplexing^9^ facilitated combined imaging of proteins and nucleic acids, while astigmatism-based axial encoding^84^ enabled imaging in 3D.

TkPAINT holds significant potential for advancing multiplexing strategies for spatial proteomics with DNA-PAINT. Current sequential DNA-PAINT schemes have, achieved up to 30-plex imaging^10,11^. Single antibody resolution in tkPAINT could enable incorporation of barcoding^87–89^ or in situ sequencing^90–92^ approaches, potentially scaling to hundreds of targets in fewer rounds. Computational methods^93,94^ and isotropic 3D imaging^95–98^ could further refine axial encoding. Additionally, tkPAINT could be combined with RESI^14^ to reach Ångstrom resolution or complement nanoscopy approaches such as MINFLUX^99^, particularly for densely-packed targets. Finally, parallel sample preparation could offer unique opportunities for correlative super-resolution and electron microscopy^34,40,100,101^, further broadening tkPAINT’s versatility.

Limitations of our study include the steric hindrance and variability in specificity inherent to antibody labeling. Smaller, stoichiometric labels, such as nanobodies^102^, genetic tags^103^, or unnatural amino acids^104^, could address these challenges, improving both structural resolution and molecular counting. High-pressure freezing and freeze substitution^105,106^ offer a promising route to further minimize fixation artifacts and capture molecular organization closer to the *in vivo* state^20,49^. The reduced imaging volume of tkPAINT compared to whole-cell imaging limits visualization of low-abundance targets and larger structures such as entire genomic regions^54^. Furthermore, compartments such as larger nuclear speckles might appear as multiple smaller structures in a single section. Serial cryosectioning^32,40,107^ could address this but would require optimization in order to mitigate challenges such as partial sample loss and folding during manual handling. Combining ultramicrotomy with resin embedding as in array tomography^45,57^ may provide an alternative, though at reduced antigenicity^35^. Implementation of machine learning^48,108,109^ and automated imager exchange^44^ could accelerate tkPAINT imaging to promote volumetric reconstructions. Nevertheless, the wealth of information gained from super-resolution studies that are based on imaging single nuclear ‘optical sections’ highlight the strong potential for studying molecular principles of genome organization in single sections alone^51–53,110–115^.

Our work enhances the potential of DNA-PAINT for single-protein imaging in various aspects. Through sectioning, we decoupled imaging performance from target selection, achieving optimal conditions for probing nanoscale organization even in dense intracellular environments. Unlike whole-cell nuclear immunolabeling, which requires disruptive permeabilization^19^, tkPAINT leverages permeabilization-free access to the nucleus in ultrastructure-preserved cells. This unique feature could enable functional studies linking nuclear and cytoplasmic mechanisms. The integration of in situ hybridization with immunolabeling extends this potential for multimodal investigations of protein–nucleic acid interactions. Finally, consistent imaging performance in both cultured cells and tissues demonstrates tkPAINT’s potential for comparative studies between cultured cells and tissues. In conclusion, we believe tkPAINT’s broad applicability will help drive DNA-PAINT toward becoming a routine tool for biological discovery.

## Extended data

**Extended Data Fig. 1.**
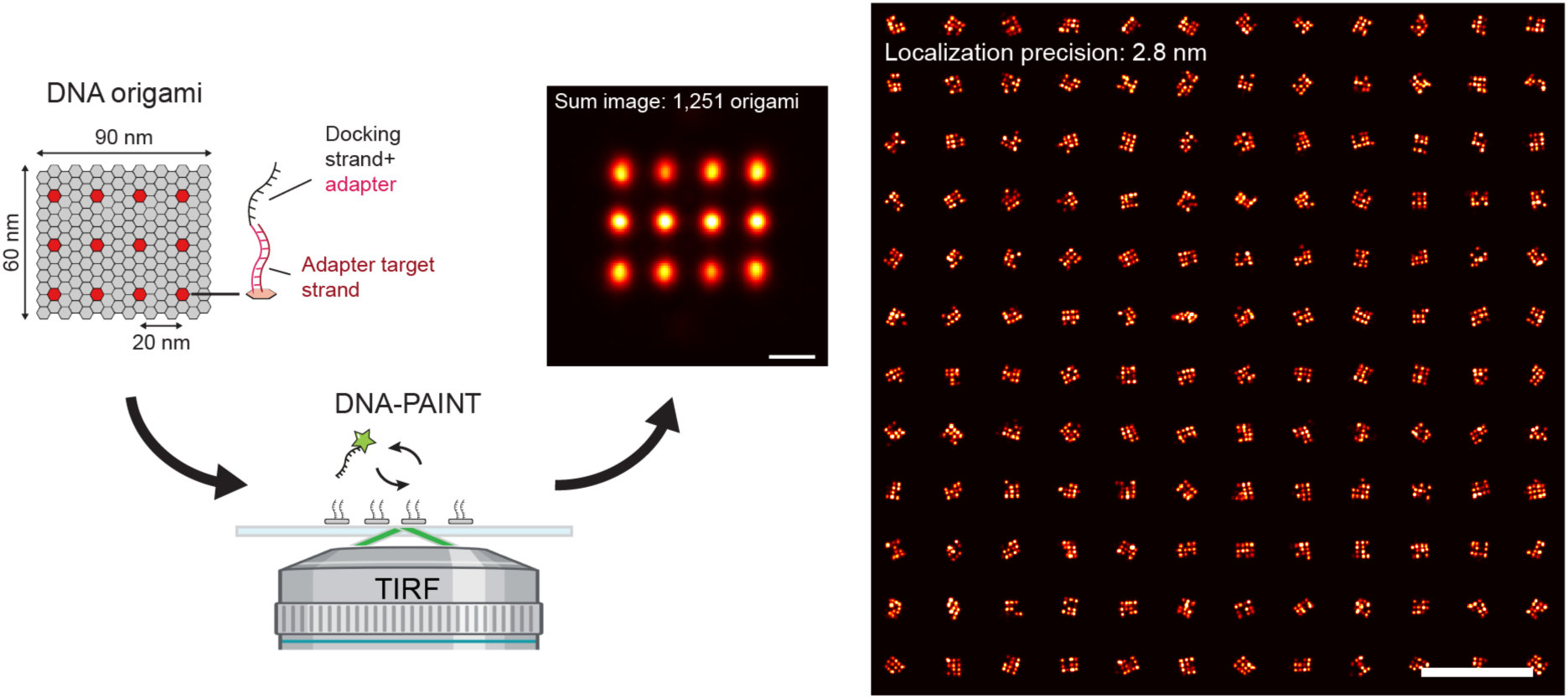
TIRF-based DNA-PAINT imaging of synthetic DNA origami 20 nm grids. DNA-PAINT image of surface-immobilized DNA origami featuring a pattern of docking strands at 20-nm spacing (’20 nm grids’^8^) acquired on our TIRF system. The origami is designed in a modular fashion by carrying 12 anchored 20nt adapter target strands (dark red) to which docking strand-adapter hybrid oligos can be stably hybridized. This way the same origami structure can be used to test different imager-docking strand combinations (see **Methods**). The increased distance between docking strand and anchor point on the origami does not lead to a noticeable decrease in resolution since the hybridized docking strand is still able to rapidly rotate around the anchor point such that on average emitted photons still allow to precisely pinpoint the anchor point^116^. Note that for space reasons some origami illustrations within this work do not show the adapter explicitly, but this origami design was exclusively used for all DNA origami experiments shown. The left image displays an averaged sum image of 1,251 origami and the right image a random selection of 144 origami arranged in 12×12 square. The localization precision for the data set is stated in the right image. Scale bars, 20 nm in left image and 200 nm in right image.

**Extended Data Fig. 2.**
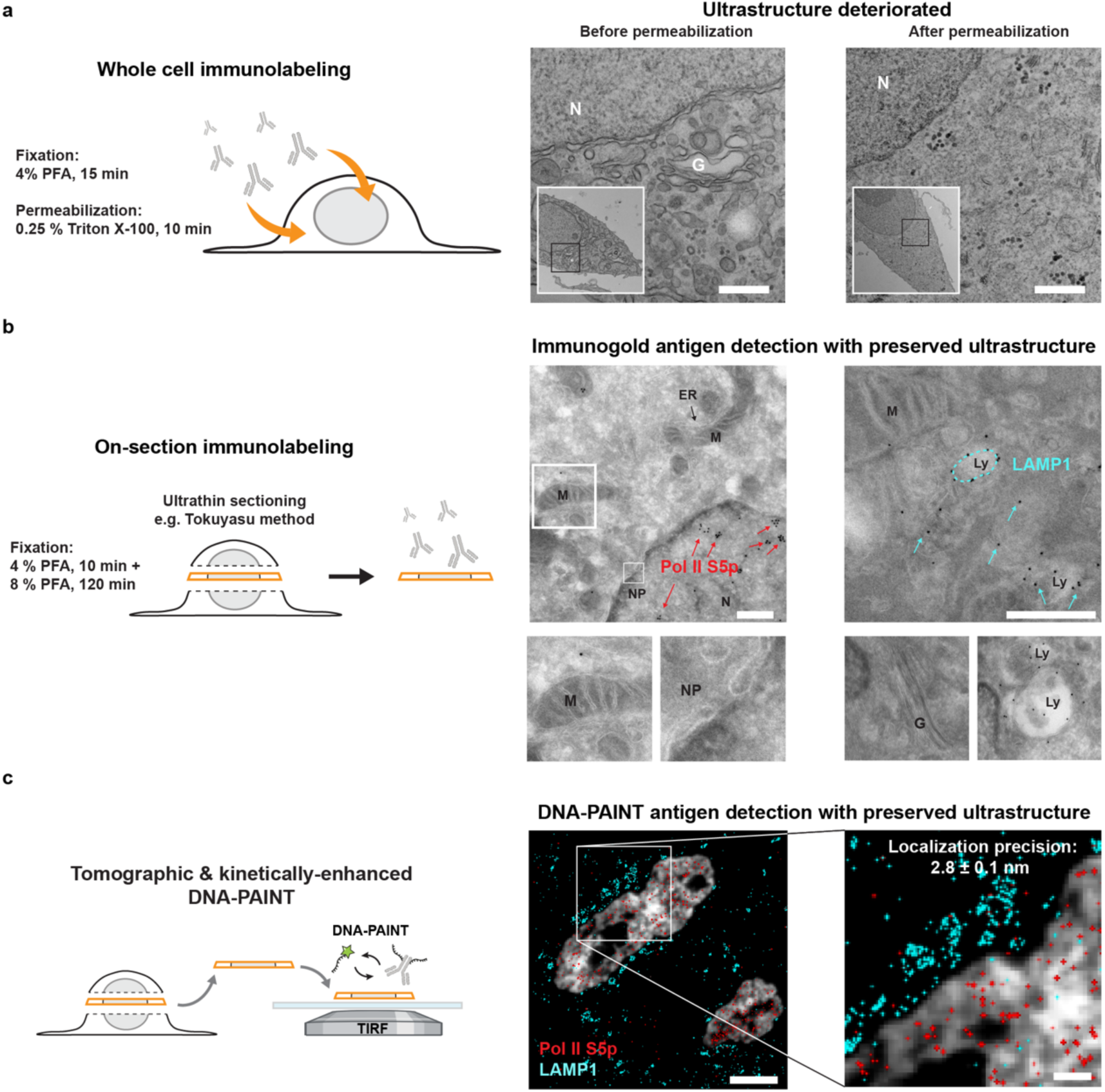
tkPAINT enables cell-wide DNA-PAINT imaging under ultrastructure-preserved conditions at sub-3 nm localization precision. **a** Ultrastructural disruption in standard paraformaldehyde(PFA)-based immunofluorescence protocols for whole cells^18^. Transmission electron microscopy (TEM) images of HeLa cells depict how brief fixation times lead to poor structural integrity and permeabilization with Triton X-100 causes reductions in cytoplasmic density as well as apparent organelle loss. Nuclear ultrastructure is relatively well-preserved even for brief fixation and permeabilization (Supplementary Fig. 4). **b** Physical sectioning enables “on-section” labeling of intracellular antigens without permeabilization. TEM images show Tokuyasu cryosections of HeLa cells prepared following a PFA-based fixation protocol optimized for ultrastructural preservation. Immunogold reveals sites of cytoplasmic LAMP1 and nuclear Pol II S5p; Mitochondria (M), nuclear pores (NP), Endoplasmic Reticulum (ER). Golgi and lysosomes (Ly) are highlighted. **c** tkPAINT principle: physical sectioning (e.g. using the Tokuyasu method) enables TIRF-based DNA-PAINT imaging of ultrastructurally-preserved specimens, even without permeabilization. The localization precision of 2.8 ± 0.1 nm was measured over four independent repeats (mean and std., respectively). Scale bars, 200 nm in (a), 400 nm in (b), 2 µm in (c) and 100 nm in zoom-in.

**Extended Data Fig. 3.**
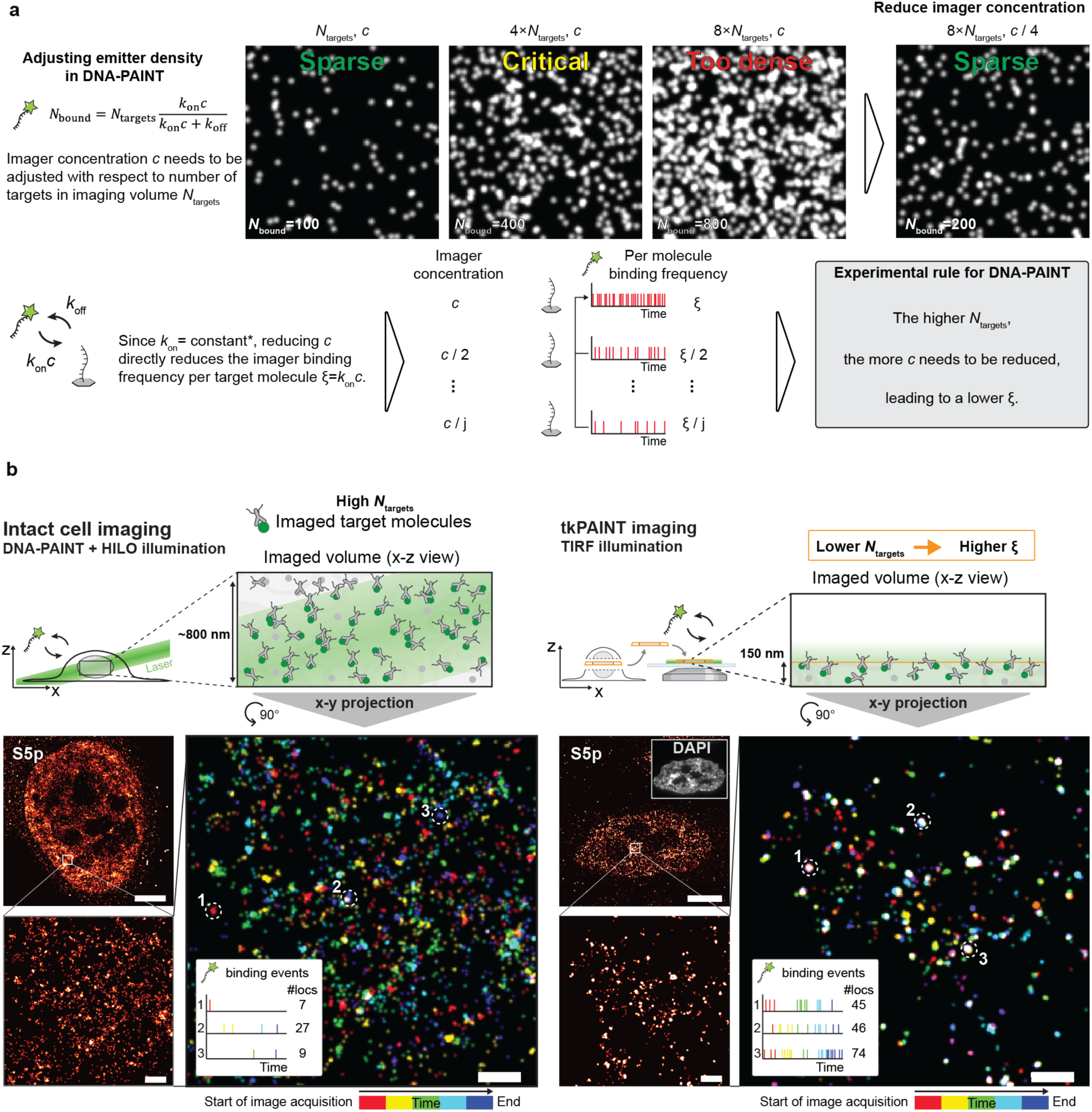
Volume reduction in tkPAINT enhances per-molecule imager binding frequencies. **a** Simulated raw DNA-PAINT images showcasing that the number of bound imagers at any given time, *N*_bound_, must be low enough to ensure sparse single-molecule blinking required for SMLM reconstruction. 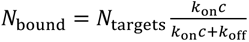, where *N*_targets_ is the number of labeled target molecules within the imaging volume, *k*_on_ is the imager association rate, *c* the imager concentration and *k*_off_ is the imager dissociation rate. *Note that *k*_on_ and *k*_off_ of a given imager-docking strand pair are constant for set experimental conditions such as temperature and buffer conditions^68^. For samples featuring a dense abundance of target molecules, *N*_bound_ can become too large and blinking events too dense, such that *c* needs to be reduced. However, a reduction in *c* inevitably leads to a decrease in the per-molecule imager binding frequency *ξ* = *k*_on_*c*. The number of randomly distributed emitters is stated in the bottom left corner of each simulated image and the image dimension are 16.64 µm x 16.64 µm. **b** Same HILO DNA-PAINT image and tkPAINT image as shown in **Fig. 1**c and d, respectively. Localizations in the zoom-in were color-coded according to registration time during data acquisition (five colors, e.g. red for first and blue for last temporal segment; total imaging time: ∼17 min. The HILO DNA-PAINT image displays largely discretely colored localizations, which is expected since the large axial imaging volume in HILO bears a large *N*_targets_, requiring to image at low imager binding frequency *ξ* that is not sufficient to repeatedly sample targeted molecules with imager binding events within the time of image acquisition. The tkPAINT image, in contrast, features concentrated accumulations of localizations of which many are revealed by temporal coloring as repetitive ‘white-colored’ localization clusters. The volume reduction in tkPAINT can thus be an effective way of enhancing imager binding statistics by reducing *N*_targets_ and thus allowing to image at higher per-molecule imager binding frequencies. For both images, time traces of imager binding and number of localizations are shown for three regions indicated by white circles, indicating high per molecule binding. Scale bars, 5 µm in (**b**) and 400 nm in zoom-ins.

**Extended Data Fig. 4.**
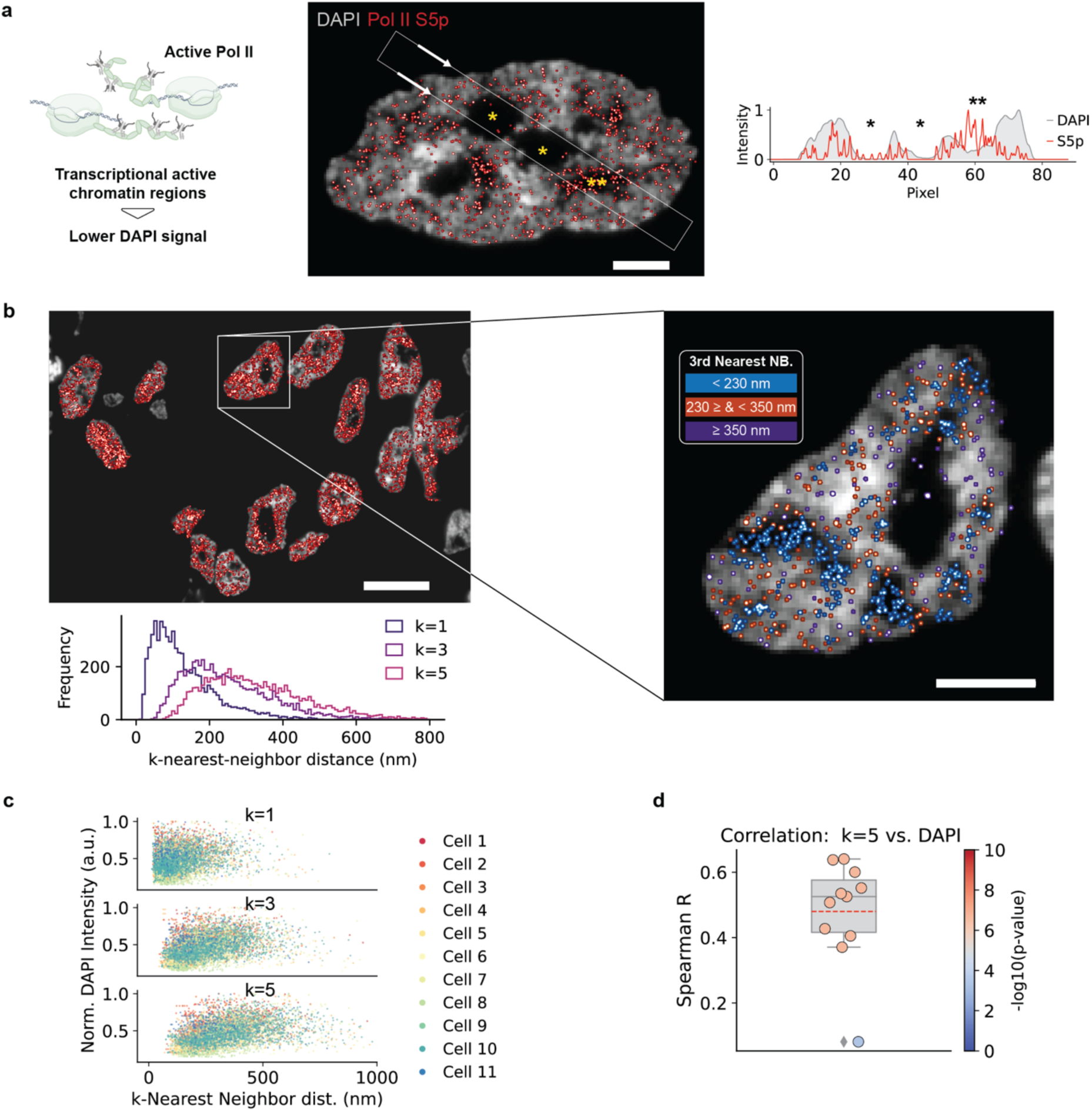
Pol II S5p correlation analysis DAPI vs. nearest neighbor distances. **a**. Left: active Pol II associates with euchromatin featuring lower DAPI intensities compared to A-T rich and densely-packed heterochromatin. Center: Pol II S5p tkPAINT image including DAPI overlay (same as shown in **Fig. 2**a). Right: The line profiles below the image show the Pol II S5p and DAPI signal distribution across the white box indicated in the center image. The box first crosses two DAPI-negative regions without Pol II S5p signal (presumably nucleoli) followed by a third DAPI-negative region featuring high S5p signal. **b** Histogram of k-nearest-neighbor distances (k=1,3,5) between Pol II S5p localization clouds for the data set shown in **Fig. 3**a. In the magnified cell on the right localization clouds were colored according to their 3^rd^ nearest neighbor distance (blue: <230 nm, red ≥ 230 nm & < 350 nm; purple ≥ 350 nm). The rendering visually confirms that DAPI-weak regions feature higher local abundances of Pol II S5p localization clouds^49,59^. Highly clustered Pol II (blue localization clouds) in DAPI weak areas likely correspond to nuclear speckles. **c** To quantify the anti-correlation between DAPI intensity (as a degree of chromatin compactness) and Pol II S5p abundance, we plotted k-nearest neighbor distances vs. normalized DAPI intensity for all nuclei in the data set. Indeed, localization clouds with small k-nearest neighbor distances, indicating a high local abundance of the antigen, were associated with DAPI weaker regions. The correlation becomes more pronounced for higher order nearest neighbor distances. **d** We calculated the Spearman rank-order correlation coefficient (R) for each cell in the data set (n=11) for the 5^th^ nearest neighbor distance of each localization cloud, confirming the correlation as highly significant (R≈0.5; p<0.0001). A correlation of <1 is expected since Pol II is absent from nucleoli, which are also DAPI weak nuclear regions. Scale bars, 10 µm in (b) and 3 µm in zoom in (b).

**Extended Data Fig. 5.**
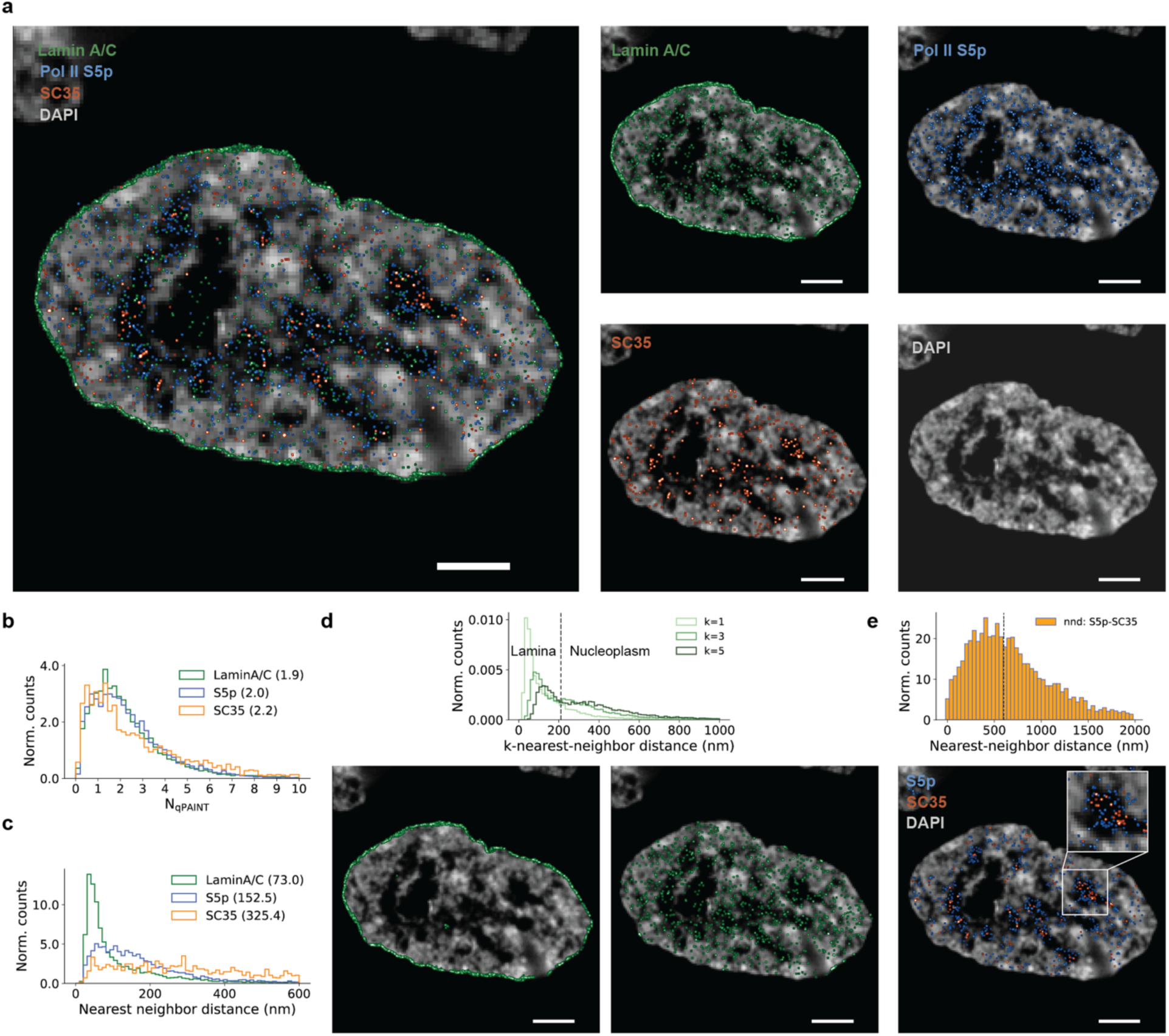
Quantitative analysis of Exchange-tkPAINT: Lamin A/C, Pol II S5p, SC35. **a** Exchange-tkPAINT image of Lamin A/C, Pol II S5p and SC35 including DAPI image (same as in **Fig. 4**a), overlayed on the left and displayed individually on the right. **b** Histogram of qPAINT counting results obtained for Lamin A/C, Pol II S5p and SC35. **c** Histogram of nearest neighbor distances measured individually for Lamin A/C, Pol II S5p and SC35. **d** Top: inspection of higher order nearest neighbor distance histograms revealed two peaks, indicating the lamina-associated fraction and the nucleoplasmic fraction of Lamin A/C, as confirmed when filtering for each peak (black dashed line) and visualizing the spatial distribution in the images below. **e** Intermolecular nearest neighbor distance measurements between Pol II and SC35. Although both antigens are known to associate with nuclear speckles^59^, the peak around 500 nm indicates a spatial segregation. Visualization of only Pol II S5p and SC35 localization clouds with a intermolecular distance <600 nm (black dashed line) in fact revealed a more centered organization of SC35 in DAPI-negative regions with S5p in the periphery, which has been similarly observed via genomics-based approaches^81^. Scale bars, 3 m in (a) and 1 m in zoom-in.

## Online Methods

### Materials

Unmodified, dye-labeled, and modified DNA oligonucleotides were purchased from Integrated DNA Technologies, Metabion and Biomers. Unmodified oligos were purified via standard desalting and modified oligos via HPLC. DNA scaffold strands were purchased from Tilibit (p7249, identical to M13mp18). Sample chambers were ordered from Ibidi GmbH (8-well 80827 and 18-well 81817). Tris 1M pH 8.0 (AM9856), EDTA 0.5M pH 8.0 (AM9261), Magnesium 1M (AM9530G) and Sodium Chloride 5M (AM9759) were ordered from Ambion. Streptavidin (S-888) Ultrapure water (15568025), PBS (20012050), 4’,6-Diamidino-2-Phenylindole, Dihydrochloride (D1306) (A39255), BSA (AM2616) and TetraSpeck™ Microspheres 0.1 µm (T7279), DMEM (10569) and Dithiothreitol (DTT) were purchased from Thermo Fisher Scientific. BSA-Biotin (A8549), Tween-20 (P9416-50ML), Glycerol (cat. 65516-500ml), (+-)-6-Hydroxy-2,5,7,8-tetra-methylchromane-2-carboxylic acid (Trolox) (238813-5G), methanol (32213-2.5L), 3,4-dihydroxybenzoic acid (PCA) (37580-25G-F), protocatechuate 3,4-dioxygenase pseudomonas (PCD) (P8279-25UN), cell scrapers (CLS353085), Triton-X 100 (93443), Gelatin from cold fish skin (G7041-500G), Formamide (F9037), RNAse A (EN0531), Sodium Azide (S2002), HEPES (H4034-100G), FastAP Alkaline Phosphatase (EF0651), Methyl cellulose 25 CP (M6385-100G), Glycine (G8898), Sodium hydroxide (P3911-1kg), methyl cellulose (M6385), dextran sulfate (D4911) 20xSSC buffer (S6639)and sucrose (S0389) was purchased from Sigma-Aldrich. 10% fetal bovine serum was purchased from Genesee Scientific (25-514). EM grade paraformaldehyde (PFA) was purchased from Electron Microscopy Services (15714). 90 nm gold nanoparticles (G-90-20-10 OD10) were purchased from Cytodiagnostics. Primary anti-Lamin A/C (mouse, 34698), anti-LAMP1 (rabbit, 9091BF) and anti a-tubulin (rabbit, 2125BF) antibodies were purchased from Cell Signaling (mouse, 34698). Primary anti-Pol II CTD S5p (rabbit, ab5131), anti-Digoxigenin (mouse, ab420) and anti-SC35 (mouse, ab11826) antibodies were purchased from Abcam. Primary anti-Pol II CTD (mouse, CTD4H8) antibody was purchased from BioLegend. Secondary donkey anti-rabbit labeled with Alexa488 was purchased from Thermo Fisher Scientific (A21206). Secondary donkey anti-rabbit and (711-005-152) and goat anti-mouse (115-005-003) antibodies were purchased from Jackson ImmunoResearch Laboratories. DBCO-modified single domain antibodies against mouse IgG (N2005-DBCO) and rabbit IgG (N2405-DBCO) as well as mouse IgG multiplexing blocker(K0102-50) were purchased from NanoTag. 0.5-mL Amino Ultra Centrifugal Filters with 50 kDa and 10 kDa molecular weight cutoffs were purchased from Millipore (UFC5050 and UFC5010, respectively). DBCO-sulfo-NHS ester cross-linker was purchased from Vector Laboratories (CCT-A124). Qubit Protein Assay (Q33211), NuPage 4-12% Bis-Tris protein gels (NP0323BOX), NuPage LDS Sample Buffer (NP0007) was purchased from Invitrogen. InstantBlue Coomassie Protein Stain was purchased from Abcam (ab119211).

### Buffers

Four buffers were used for sample preparation and imaging: Buffer A (10 mM Tris-HCl pH 7.5, 100 mM NaCl); Buffer B (5 mM Tris-HCl pH 8.0, 10 mM MgCl2, 1 mM EDTA); Buffer C (1× PBS, 500 mM NaCl); 10x folding buffer (100 mM Tris,10 mM EDTA pH 8.0, 125 mM MgCl2). Antibody storage buffer: 1% BSA, 0.1% Sodium Azide, 10 mM EDTA, 50% glycerol). Buffers were checked for pH. Imaging buffers were supplemented with oxygen scavenging & triplet state quenching system 1× PCA, 1× PCD, 1× Trolox prior to imaging.

### PCA, PCD, Trolox

100× Trolox: 100 mg Trolox, 430 μL 100% Methanol, 345 μL 1 M NaOH in 3.2 mL H2O. 40× PCA: 154 mg PCA was mixed with 10 mL water adjusted to pH 9.0 with NaOH. 100× PCD: 9.3 mg PCD, 13.3 mL of buffer (100 mM Tris-HCl pH 8, 50 mM KCl, 1 mM EDTA, 50% glycerol).

### DNA origami design and assembly

DNA origami with 20-nm spaced docking strands (’20 nm grids’) were designed previously using the Picasso Design^8^ module. A list of all used DNA strands can be found in ref.^117^. Folding of structures was performed using the following components: single-stranded DNA scaffold (0.01 µM), core staples (0.1 µM), biotin staples (0.01 µM), extended staples for DNA-PAINT (each 1 µM), 1x folding buffer in a total of 50 µl for each sample. Annealing was done by cooling the mixture from 80 °C to 25 °C in 3 hours in a thermocycler. Using a 1:1 ratio between scaffold and biotin staples allows sample preparation without prior DNA origami purification, where otherwise free biotinylated staples would saturate the streptavidin surface and prevent origami immobilization on the glass surface. As docking strand sequence, we used a 20nt adapter motif^16^ (A20: AAGAAAGAAAAGAAGAAAAG), which allowed us to later hybridize any desired docking strand imaging to the origami via a stably-binding complementary adapter ‘cA20_DS’. The adapter motif is cA20: CTTTTCTTCTTTTCTTTCTT which is concatenated to the docking strand of choice DS (see **Supplementary Table 2** for sequences).

### DNA origami sample preparation

Ibidi 8-well slides were prepared as follows. A 10 µl drop of biotin-labeled bovine albumin (1 mg/ml, dissolved in buffer A) was placed at the chamber center and incubated for 2 min and aspirated. The chamber was then washed with 200 µl of buffer A, aspirated, and then a 10 µl drop streptavidin (0.5 mg/ml, dissolved in buffer A) was placed at the chamber center and incubated for 2 min. After aspirating and washing with 200 µl of buffer A and subsequently with 200 µl of buffer B, a 10 µl of DNA origami (1:100-200 dilution in buffer B from folded stock) was placed at the chamber center and incubated for 5 min. Next, the chamber was washed with 200 µl of Buffer B and docking strand adapters hybridizing to the DNA origami were added at 100 nM in Buffer B, incubated for 5 min and washed with 200 µl Buffer B. Finally, Buffer C and imager strand was added for DNA-PAINT imaging.

### Conjugation of secondary antibodies/nanobodies with docking strands

DNA antibody conjugations were performed in 0.5-mL Amino Ultra Centrifugal Filters with 50 kDa molecular weight cutoffs with DBCO-sulfo-NHS ester cross-linker, which was dissolved at 20 mM DMSO and stored in single-use aliquots at −80° C. This cross-linker links azide-functionalized DNA oligonucleotides to surface-exposed lysine residues. Azide-functionalized DNA oligonucleotides were stored in 1 mM deionized water. Critically, all antibodies were ordered carrier-free, as common preservatives such as bovine serum albumin and sodium azide interfere with the conjugation reaction. First, 500 µL PBS was added to the Amicon filters, which were centrifuged for 5 min at 10,000 rcf. After wetting the filters, 25 µg antibody was added and washed twice with PBS. For each wash, PBS was added to a total volume of 500 µL, and the filters were centrifuged for 5 min at 10,000 rcf. If after the second spin, the total volume remaining in each filter was greater than 100 µL, the filters were centrifuged again for 5 min at 10,000 rcf. After the second PBS wash, a 20-fold molar excess of DBCO-sulfo-NHS ester cross-linker and a 20-fold molar excess of DNA oligonucleotide were added, and after gentle mixing, each conjugation reaction was incubated in the dark at 4° C overnight. The following day, conjugated antibodies were washed three times with PBS, as described above. To elute the antibody, the filter was inverted in a fresh tube and centrifuged for 2 min at 1,500 rcf. The conjugated antibody was transferred to a clean tube and stored at −20° C in antibody storage buffer. Concentrations were measured using the Qubit Protein Assay. DNA-antibody conjugation was confirmed by comparing unconjugated and conjugated antibodies on NuPage 4-12% Bis-Tris protein gels. For each sample, 0.5 µg total protein was added to NuPage LDS Sample Buffer and 50 mM DTT. Protein was denatured at 80° C for 10 min. Gels were run at 75 V for 5 min, then at 180 V for 60 min. Gels were stained with InstantBlue Coomassie Protein Stain for 15 minutes at room temperature, rinsed with water, and imaged on a Sapphire Biomolecular Imager (Azure Biosystems).

Conjugation of DBCO-modified nanobodies (also “single domain antibody”) was performed analogously, but in 0.5-mL Amino Ultra Centrifugal Filters with 50 kDa molecular weight cutoffs. After filter wetting and washing, 25 µg nanobody was added and washed twice. For each wash, PBS was added to a total volume of 500 µL, and the filters were centrifuged for 5 min at 10,000 rcf. After the second PBS wash, a 5-fold molar excess of DNA oligonucleotide were added, and after gentle mixing, each conjugation reaction was incubated in the dark at 4° C overnight. The next day, conjugated nanobodies were washed three times and transferred to a clean tube for storage at −20° C in antibody storage buffer. Concentrations were measured using the Qubit Protein Assay and working aliquots were adjusted to 5 mM in antibody storage buffer as recommended by the manufacturer.

### Tissues

Mouse tissue was obtained from naïve control mice meeting experimental endpoint on an approved Harvard Medical School/Longwood Medical Area IACUC protocol.

### Cell culture and plating

HeLa cells were maintained in DMEM supplemented with 10% fetal bovine serum at 37 °C with 5% CO2 and were checked regularly for mycoplasma contamination. For imaging of whole HeLa cells, ∼16K cells were seeded in each well of an Ibidi 18-well chamber, placed in the incubator overnight and fixed the following day. For preparation of cell pellets for cryosectioning, ∼1 million cells were seeded in 10-cm dishes and placed in the incubator until reaching 70 % confluency.

### HeLa cell preparation for cryosectioning

HeLa cells were processed according to previously published protocols^64^. In brief, HeLa cells were grown in 10 cm Petri dishes and once reaching 70 % confluence, were fixed in 4% PFA 250 mM HEPES, pH 7.6 for 10 min. Fixative was decanted and cells further fixed with 8% PFA in 250 mM HEPES, pH 7.6 for a total of 2h at 4°C. During fixation, cells were gently scraped off the surface unidirectionally using cell scrapers previously soaked in fixative to avoid sticking. Detached cell suspension was transferred into a 1.5 mL hydrophobic Eppendorf tube and centrifuged at increasing speeds to form a pellet of fixed cells: 300 × g, 5 min; 500 × g, 2 min; 1,000 × g, 2 min; 2,000 × g, 2 min; 4,000 × g. At this point, the pellet could be resuspended in 1% PFA in 250 mM HEPES, pH 7.6 and stored overnight at 4°C. Next, the pellet was transferred between several drops of 2.1 M sucrose drops to wash away residual fixative and infiltrated 2-4h in 2.1 M sucrose (sucrose acts as cryoprotectant to prevent structural damage during freezing. The pellet becomes transparent). Next, the infiltrated pellet was transferred to a metal pin, residual sucrose carefully removed using filter paper and the pellet shaped into a cone under a dissecting light microscope and using forceps. Finally, the cell pellet was frozen by immersion into liquid nitrogen and was stored indefinitely in liquid nitrogen tanks. We would like to also highlight alternative protocols based on gelatin embedding, which can improve probe handling as discussed in a recent review^35^.

### Tissue preparation for cryosectioning

Mouse cerebellum and spleen were sectioned into 1-2 mm cubes and incubated consecutively in 4% PFA 250 mM HEPES, pH 7.6, in 8% PFA in HEPES for 2 hours at 4°C, and in 1% PFA in HEPES overnight at 4°C. Tissue cubes were then embedded in 7.5% gelatin, 10% Sucrose in PBS (gelatin-sucrose solution was prepared at 70°C and stored in 10mL aliquots at −20°C). Tissues were infiltrated in liquid gelatin-sucrose for 30 minutes at 37°C and subsequently solidified at 4°C. Then, the gel block was removed from the tube, the tissue block cut out as 1mm blocks and transferred into 2.1 M sucrose in PBS for 4h. Lastly, sucrose-infiltrated tissue blocks were placed on metal pins, residual sucrose carefully removed using filter paper, frozen by immersion into liquid nitrogen and stored indefinitely in liquid nitrogen tanks.

### Tokuyasu cryosectioning

All Tokuyasu cryosectioning was performed at the Harvard Electron Microscopy Core using a Leica EM UC7 Ultramicrotome equipped with a FC7 cryo-chamber. Frozen cell/tissue samples were cut at a temperature of −110°C using a diamond knife (Diatome). Lastly, sections were collected using drops a freshly prepared 1:1 mixture of 2.1 M sucrose in PBS and 2% methyl cellulose in water and transferred onto Ibidi 8-well chambers for tkPAINT imaging, that had previously been glow discharged (EMS100x, 2min at 40mA). Sectioned samples can be stored at −20°C for months.

### TEM imaging of Tokuyasu sections

For transmission electron microscopy imaging, cryosections were placed on formvar-coated grids, washed, and contrasted using methyl cellulose/uranyl acetate. TEM imaging was performed at the Harvard Electron Microscopy Core on a JEOL 1200EX TEM.

### HeLa cell fixation, epon embedding and sectioning for TEM imaging

HeLa cells were grown in 10 cm Petri dishes and once reaching 70 % confluence, were fixed in 4% PFA 250 mM HEPES, pH 7.6 for 15 min followed by three washes in PBS for 2 min each. For permeabilized samples, permeabilization was applied in Petri dish, followed by three washes in PBS. Cells were gently scraped and collected into a 0.5 mL tube and centrifuged at 200 × g for two min to form pellets. The solution was exchanged and pellets stored in 1% PFA in 250 mM HEPES, pH 7.6 overnight at 4 °C. The next day, cell pellets were postfixed with 1% Osmium Tetroxide (OsO4)/1.5% Potassium Ferrocyanide(KFeCN6) for 1 hour, washed 2× in water, 1× Maleate buffer (MB) 1× and incubated in 1 % uranyl acetate in MB for 1 hr followed by 2 washes in water and subsequent dehydration in grades of alcohol (10 min each; 50%, 70%, 90%, 2×10min 100%). The samples were then put in propylene oxide for 1 h and infiltrated ON in a 1:1 mixture of propylene oxide and TAAB Epon (TAAB Laboratories Equipment Ltd, https://taab.co.uk). The following day the samples were embedded in TAAB Epon and polymerized at 60 °C for 48 h. Ultrathin sections (∼80 nm) were cut on a Reichert Ultracut-S microtome, picked up onto copper grids, stained with lead citrate and examined in a Tecnai Spirit BioTwin Transmission electron microscope. Images were recorded with an AMT NanoSprint43-MkII camera.

### Immunogold TEM imaging

Tokuyasu sectioning was performed at −120 °C and at ∼80nm cryosection thickness. Sections were picked up on a drop of 2.3 M sucrose with a small amount of 2% methyl cellulose added (9:1 mixture) and transferred to formvar-carbon coated copper grids. Gold labeling was carried out at room temperature on a piece of parafilm: antibodies were diluted in 1% BSA in PBS Grids, floated on drops of 1% BSA for 10 minutes to block for unspecific labeling, transferred to 5 µl drops of primary antibody and incubated for 30 minutes. Subsequently, grids were washed in 4 drops of PBS (total 10 min) before incubation in 10nm Protein A-gold (University Medical Center, Utrecht, the Netherlands) for 20 min. Grids were washed in 2 drops of PBS followed by 4 drops of water (total 15 min). The labeled sections were contrasted and embedded in methyl cellulose by floating the grids on a mixture of 0.3% uranyl acetate in 2% methyl cellulose for 5 minutes before blotting excess liquid off on a filter paper. Grids were imaged on a JEOL 1200EX Transmission electron microscope and an AMT 2k CCD camera.

### Labeling of cryosections for tkPAINT

8-well chambers containing cryosections were thawed and washed 3× in PBS under agitation for 10min for sucrose removal and quenched with 100mM glycine in 100mM HEPES for 15min. Next, cryosections were permeabilized in 0.3% Triton-X 100 in PBS for 5min, rinsed 3× in PBS and ready for subsequent labeling. Note, that Tokuyasu immunogold protocols vary regarding antibody incubation times. A general rule of thumb is using high antibody concentrations and short incubation times, rather than low concentrations for extended incubations^17^. Hence, we chose relatively high antibody dilutions (1:50-200) and could even observe strong antibody signal for incubations as short as 5min. For a systematic investigation, antibody titration series can be advised. For our proof-of-concept study we applied varying blocking and/or labeling conditions, which are listed in **Supplementary Table 1** for all experiments with respect to blocking buffer as well as both antibody dilution and incubation times. The blocking buffer was used for both antibody/nanobody incubations and as a washing solution in between labeling in case of indirect primary and secondary antibody/nanobody labeling. After antibody labeling, cryosections were washed 3× in PBS, stained with 30 nM DAPI in PBS for 3min and washed again with PBS. For all tkPAINT experiments based on secondary nanobodies the samples were postfixed in 4% PFA for 5 min, followed by three washes in PBS prior to imaging and DAPI staining. Lastly, Buffer C and imager was added for tkPAINT imaging. Note that DAPI staining could faint for several rounds of washing, especially for Exchange-PAINT experiments. However, staining could be simply recovered by performing another round of DAPI staining at the same concentration as stated above.

#### Phosphatase control (Fig. 2c)

Two cryosection samples were processed as previously described until the blocking step, at which they were placed for 1h at 37°C and one incubating with alkaline phosphatase to remove phosphorylation site S5p as target antigen^59^. After washing 3× in PBS, normal blocking and indirect immunostaining was performed using a fluorescently-labeled secondary antibody.

#### Combined a-tubulin and telomere imaging (Fig. 4c)

Cryosections were labeled for a-tubulin using primary antibody + secondary antibody incubation and postfixed with 4% PFA in PBS for 10min followed by a 10min glycine quenching step. Next, the samples were washed with PBS, and incubated with 100-fold diluted RNase A/T1 Mix in 1× PBS at 37 °C for 1 h. Samples were washed 3× in PBS, rinsed and incubated with 50% formamide in 2× SSC for 15min. Next, the sample was placed on a heat block at 90 °C for 4.5 min in 50% formamide in 2× SSC. A 20nt FISH probe against telomeric repeat (AACCCTAACCCTAACCCTAA-A488) was added at 1 µm concentration in 20% formamide, 10% dextran sulfate and 4× SSC and incubated overnight at 37°C for hybridization. Lastly, the sample was washed 2× with 20% formamide 2× SSC, rinsed with PBS, and 30 nM DAPI in PBS for 3min was added. After a final wash in PBS, Buffer C was added and imager for tkPAINT imaging.

#### mRNA imaging via poly(dT) probes (Fig. 4d)

Cryosections were treated as described until the blocking step, followed by a 10min wash in 4x SSC. Next, 40nt poly(dT) probe modified with digoxigenin were added in 20% hybridization buffer (20% ethylene carbonate, 10% dextran sulfate and 4× SSC) buffer at 37 °C overnight in a humidity chamber. The next day, the sample was washed 2× with 20% EC 2xSSCT for 15min, followed by three rinses with 4× SSC. The sample was then blocked with 1% gelatin in PBS for 10min and subsequently subject to indirect immunostaining as described in **Supplementary Table 1**. After final washes, Buffer C and imager was added for tkPAINT imaging.

### Fixation and labeling for whole HeLa cell imaging

24h after seeding HeLa cells in Ibidi 18-well chambers, cells were fixed using 4% PFA 250 mM HEPES, pH 7.6 for 20min. Next, samples were washed 4× in PBS (30s, 60s, 2×5 min) and both blocked and permeabilized in 3% BSA and 0.25% Triton X-100 in PBS at room temperature for 90 min. Primary rabbit anti-Pol II S5p antibody was added at 1:100 in 3% BSA and 0.1% Triton-X 100 in PBS and incubated overnight at 4 °C. The next morning, samples were washed 4x washes in PBS (30s, 60s, 2× 5min) and DNA-conjugated secondary antibody (1:100) was added at 1:100 in 3% BSA and 0.1% Triton-X 100 in PBCS and incubated for 1h at room temperature. Samples were quickly washed 3× in PBS, incubated with gold particles as fiducial markers (1:20 in PBS) for 5 min, washed again 2× in PBS before adding Buffer C and imager for DNA-PAINT imaging.

### Super-resolution microscopy setup

TIRF and HILO imaging was carried out at MicRoN Imaging Core at Harvard Medical School on a Nikon Ti inverted microscope equipped with a Nikon Ti-TIRF-EM Motorized Illuminator, a Nikon LUN-F Laser Launch with single fiber output (488nm, 90mW;561 nm, 70mW; 640nm, 65mW) and a Lumencore SpectraX LED Illumination unit. The objective-type TIRF system with an oil-immersion objective (Apo TIRF 100×/1.49 DIC N2). DNA-PAINT experiments were performed using the 560 nm laser line and fluorescence emission was passed through a Chroma ZT 405/488/561/640 multi-band pass dichroic mirror mounted on a Nikon TIRF filter cube located in the filter cube turret and a Chroma ET 595/50m band pass emission filter located on a Sutter emission filter wheel within the infinity space of the stand before image recording on a line on a sCMOS camera (Andor, Zyla 4.2) mounted to a standard Nikon camera port. For astigmatism-based 3D imaging, the C-mount side port of the microscope body was replaced by a custom-built construction allowing to insert a cylindrical lens in front of the camera (description including component list in **Supplementary Fig. 12**).

### Imaging conditions

All fluorescence microscopy data was recorded with the sCMOS camera (2048 × 2048 pixels, pixel size: 6.5 µm). Both microscope and camera were operated with the Nikon Elements software at 2×2 binning and cropped to the center 512 × 512 pixel field-of-view. The camera read out rate was set to 200 MHz and the dynamic range to 16 bit. For detailed imaging parameters specific to the data presented in all main and supplementary figures refer to **Supplementary Table 1**.

### Image analysis

Please refer to **Supplementary Fig. 2** and **Supplementary Fig. 5** for a detailed step-by-step illustration through all processing steps of super-resolution reconstruction. All DNA-PAINT/tkPAINT imaging data was processed and reconstructed using the Picasso^8^ software suite, Fiji^118,119^ and previously-published^16,116^ and custom Python modules.

## Data availability

All data are available in the main text or the supplementary materials, and materials are available upon request.

## Code availability

Super-resolution reconstruction was performed using the Picasso^8^ suite developed by the Jungmann lab: https://github.com/jungmannlab/picasso.Previously-published custom Python packages employed in this study are available in public repositories: https://github.com/schwille-paint/picasso_addon and https://github.com/schwille-paint/lbFCS2. Additional custom code will be made available upon request.

## Acknowledgements

We thank Paula Montero Llopis and Praju Vikas Anekal at HMS MicRoN Imaging facility, Margaret Coughlin and Anja Nordstrom at HMS Electron Microscopy Core, Tom Ferrante, Maurice Perez and Talley Lambert for valuable experimental support. We thank Miiko Sokka and Nicola Neretti for sharing and providing antibodies on short notice and for helpful discussions. We thank Oliver Dodd and Soufiane Aboulhouda for sharing mouse tissues. We thank Robert Tjian, Thomas Graham, Claudia Cattoglio and Nam Che for sharing cell lines and plasmids as well as Merrick Pierson Smela for help regarding plasmid recovery and transfection. We thank Silvia Filipa Carvalho and Izabela-Cezara Harabula for sharing protocols and discussing Tokuyasu cryosectioning for fluorescence microscopy. We thank the Bewersdorf lab at Yale University for sharing fluorogenic imagers and docking strand-conjugated secondary antibodies. We thank Gareth Griffiths and Heinz Schwarz for helpful discussions and invaluable expert advice with regarding the Tokuyasu-Method. We thank Nuno Martins, Fei Zhao, Antonios Lioutas, Jumana Alhaj Abed, Tae Ryu, Eunice Fabian-Morales and Erkin Kuru for manuscript feedback and helpful discussions. We further acknowledge helpful discussions with Hylkje Geertsema, Yolanda Markaki, Bas van Steensel, Merle Hantsche-Grininger, Peter Becker, Christophe Leterrier and Hiroshi Sasaki. We acknowledge biorender.com which was partially used in the illustrations. J.S. acknowledges support by the European Molecular Biology Organization (ALTF 816-2021), R.B.M. acknowledges support by NSF (Graduate Research Fellowship grant 2140743) and NIH (Molecular Biophysics Training grant NIGMS T32 GM008313). P.Y. acknowledges support by the NIH (Pioneer Award DP1GM133052). A.P. acknowledges support by the Helmholtz Association. G.M.C. acknowledges support by the Department of Energy (DE-FG02-02ER63445). C.-t.W. acknowledges funding from NIH (4D Nucleome Program 5RM1HG011016-03 and 5UM1HG011593).

## Author contributions

J.S. conceived the study, performed experiments, analyzed data and wrote the manuscript with input from all authors. M.E. performed cryosectioning, provided cryosection samples and performed TEM imaging. M.N. conjugated antibodies, maintained cell culture and provided cell samples for cryoblock preparations. L.M. prepared tissue cryoblocks and performed initial tissue tkPAINT experiments. S.A. contributed to initial joint experiments and to the manuscript storyline. R.B.M. performed antibody conjugations. L.B. developed code for chromatic aberration correction, nuclear volume calculation and contributed to the manuscript storyline. C.P.H. contributed to initial experiments. A.W. and L.A.-J. provided practical training and supported early protocol development J.W. provided helpful advice regarding cryosectioning and TEM data interpretation. P.Y. provided laboratory infrastructure, helpful advice and shared reagents. A.P. hosted a lab visit of J.S., provided training, protocols related to cryosectioning and valuable manuscript input. G.M.C. and C.-t.W. guided the project through joint discussions, valuable feedback and contributed to the manuscript storyline. All authors read and approved the final manuscript.

