## Supplementary Information for "Cryosectioning-enhanced super-resolution microscopy for single-protein imaging across cells and tissues"

#### Supplementary Figures


| | |
|---|---|
| Supplementary Fig. 1 | TEM imaging confirms nuclear preservation in permeabilized cryosections |
| Supplementary Fig. 2 | Segmentation of nuclear localizations for quantitative comparison of HILO and tkPAINT |
| Supplementary Fig. 3 | tkPAINT does not require fluorogenic imagers for bleaching protection |
| Supplementary Fig. 4 | Permeabilization enhances nuclear epitope accessibility |
| Supplementary Fig. 5 | DBSCAN localization cloud detection, quantitative analysis and kinetic filtering |
| Supplementary Fig. 6 | Kinetic filtering efficiently removes non-repetitive localization clouds and allows quantitative evaluation of labeling strategies |
| Supplementary Fig. 7 | Influence of section thickness on tkPAINT imaging |
| Supplementary Fig. 8 | Scan for convergent minimum number of localizations per cloud – HeLa cells |
| Supplementary Fig. 9 | qPAINT analysis steps for DNA origami data |
| Supplementary Fig. 10 | qPAINT analysis steps for tkPAINT data |
| Supplementary Fig. 11 | Scan for convergent minimum number of localizations per cloud – mouse tissues |
| Supplementary Fig. 12 | Custom cylindrical lens insertion for 3D tkPAINT |
| Supplementary Fig. 13 | Planar fit for axial tilt-correction in 3D tkPAINT |
| Supplementary Fig. 14 | Axial distribution of antibody signal for 3D Exchange-tkPAINT |


#### Supplementary Tables


| | |
|---|---|
| Supplementary Table 1 | Imaging parameters for tkPAINT/DNA-PAINT |
| Supplementary Table 2 | Used DNA oligonucleotide sequences as labels |


#### Supplementary References

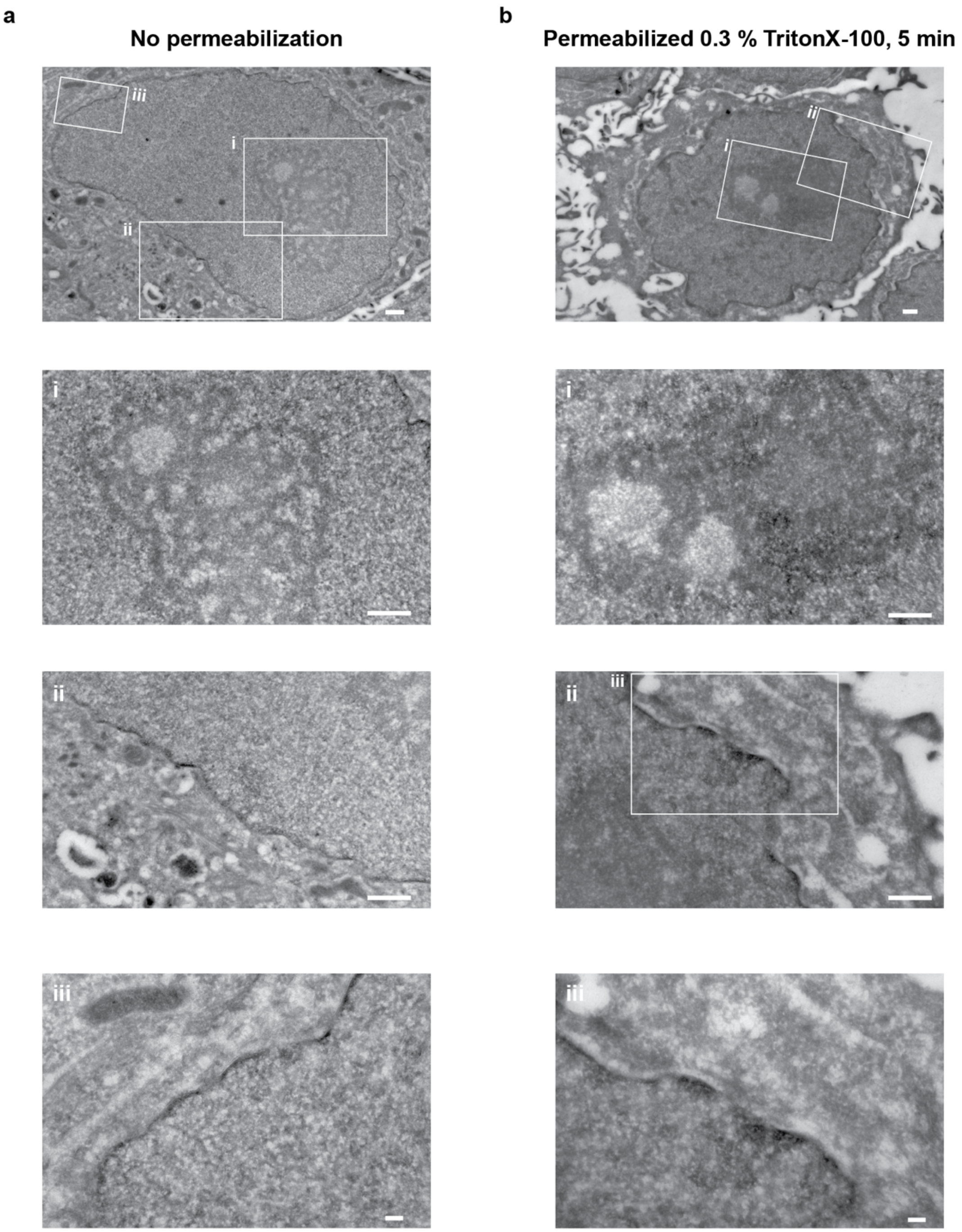


**Supplementary Fig. 1 | TEM imaging confirms nuclear preservation in permeabilized cryosections. a** Transmission electron microscopy (TEM) images of untreated 150 nm HeLa cryosections. Although ultrastructural studies are commonly performed in thinner cryosection (<100 nm), the contrast is sufficient to reveal structural features within the nucleus (nucleolus, nuclear envelope, and nuclear pores) and in the cytoplasm (endoplasmic reticulum, mitochondria, and cristae). The bottom three images show magnified regions of the same cell shown in the top image. **b** TEM images of 150 nm cryosections that were permeabilized with 0.3 % TritonX-100 for 5 min. As expected, detergent treatment affects lipids and membrane structures and results in observable extraction especially from the cytoplasmic domain. Overall, the intranuclear space, chromatin as well as the nucleolus appear ultrastructurally preserved. The bottom three images show magnified regions of the same cell shown in the top image. Scale bars, 500 nm, except 100 nm in (**a**iii and **b**iii).

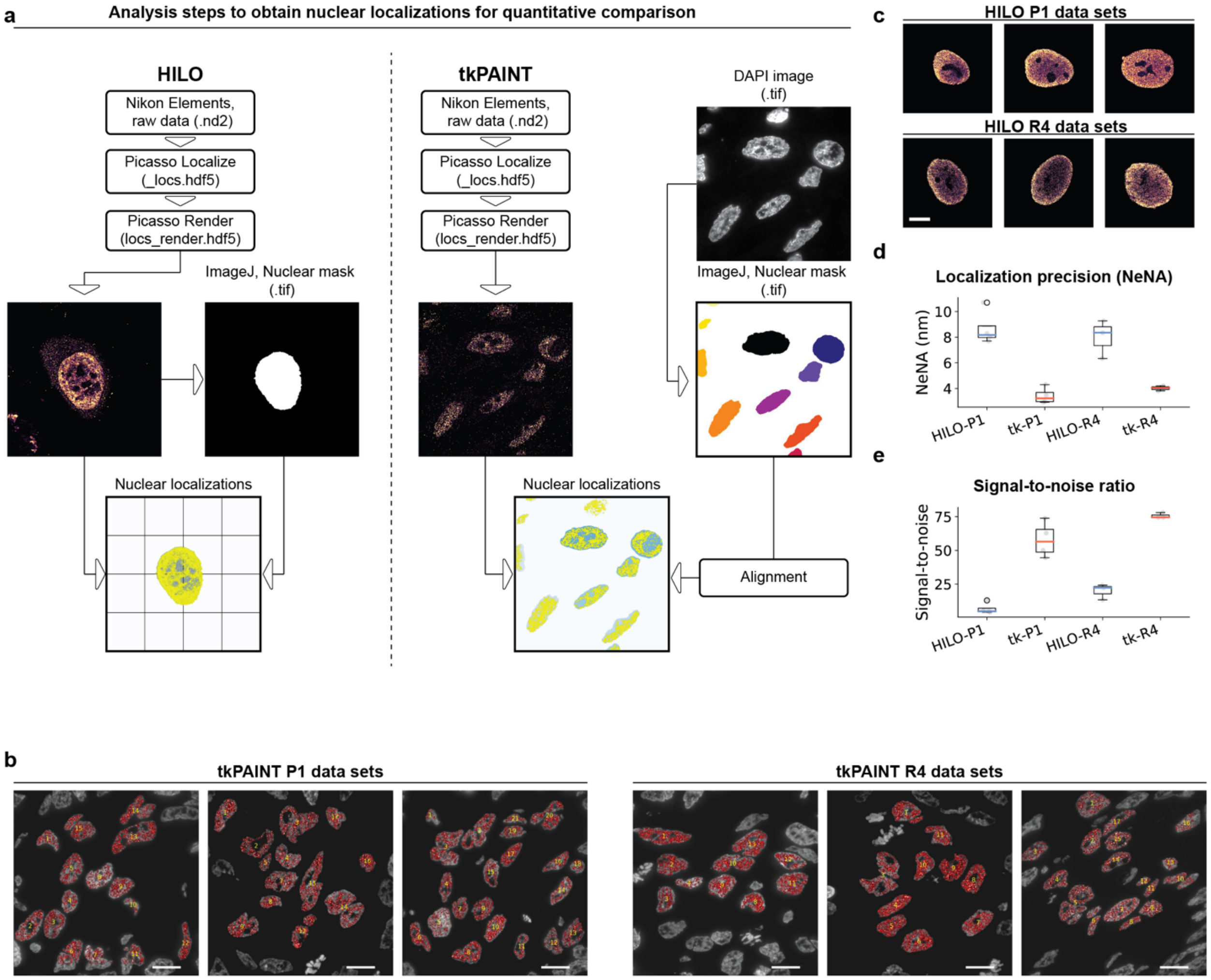


**Supplementary Fig. 2 | Segmentation of nuclear localizations for quantitative comparison of HILO and tkPAINT. a Left**: Workflow for nuclear segmentation in HILO DNA-PAINT imaging. Raw time series were recorded using Nikon Elements and processed with Picasso Localize software module[1]. The localization files were subsequently loaded into Picasso Render[1] to perform drift correction (global correction via redundant-cross correlation[2] and subsequent correction based on gold particles as fiducials) and saved. From Picasso Render we further exported a low-resolution and oversaturated image, that we used to create a nuclear mask via FIJI[3] (thresholding to create a mask & the plugin BIOP/Image Analysis/ROIs/ ROIs to label image to export a binary mask as .tif file. https://github.com/BIOP). Using a custom Python script, both the drift corrected localization file and the binary mask were loaded to filter for nuclear localizations based on the mask. Note: we did not use DAPI staining for nuclear segmentation as in tkPAINT due to poor image quality in HILO illumination. **Right**: Workflow for nuclear segmentation in tkPAINT imaging. Data acquisition, localization and drift correction were performed as in a, only that individual localization clouds could serve as fiducials for drift correction and no gold particles were needed, similar to imaging DNA origami[1]. Prior to each tkPAINT experiment, we acquired a single DAPI image for later nuclear segmentation. For each tkPAINT experiment, a corresponding binary mask was created out of the DAPI image analogously to a. We initially performed an affine transformation to match DAPI and 560 nm channel (Cy3b) using Fiji[3] for descriptor-based registration[4] based on TetraSpeck™ multicolor bead images, but found that the effect on the diffraction limited DAPI mask was negligible. Hence, a Python script was used to directly align DAPI masks and tkPAINT datasets, correcting for a potential lateral offset due to sample drift during the tkPAINT acquisition, and to subsequently filter for all nuclear localizations accordingly. **b**. tkPAINT Pol II S5p datasets acquired using standard imager sequence P1 and speed imager sequence R4. Interphase nuclei that entirely lied within the aligned DAPI mask after drift correction were selected. **c** HILO DNA-PAINT Pol II S5p datasets acquired using standard imager sequence P1 and speed imager sequence R4. **d** Localization precision (NeNA – Nearest Neighbor Analysis[5]) for tkPAINT and HILO datasets in b-c. **e** Signal-to-noise ratio for tkPAINT and HILO datasets in b-c. **f** Nuclear localization density for tkPAINT and HILO datasets in b-c.

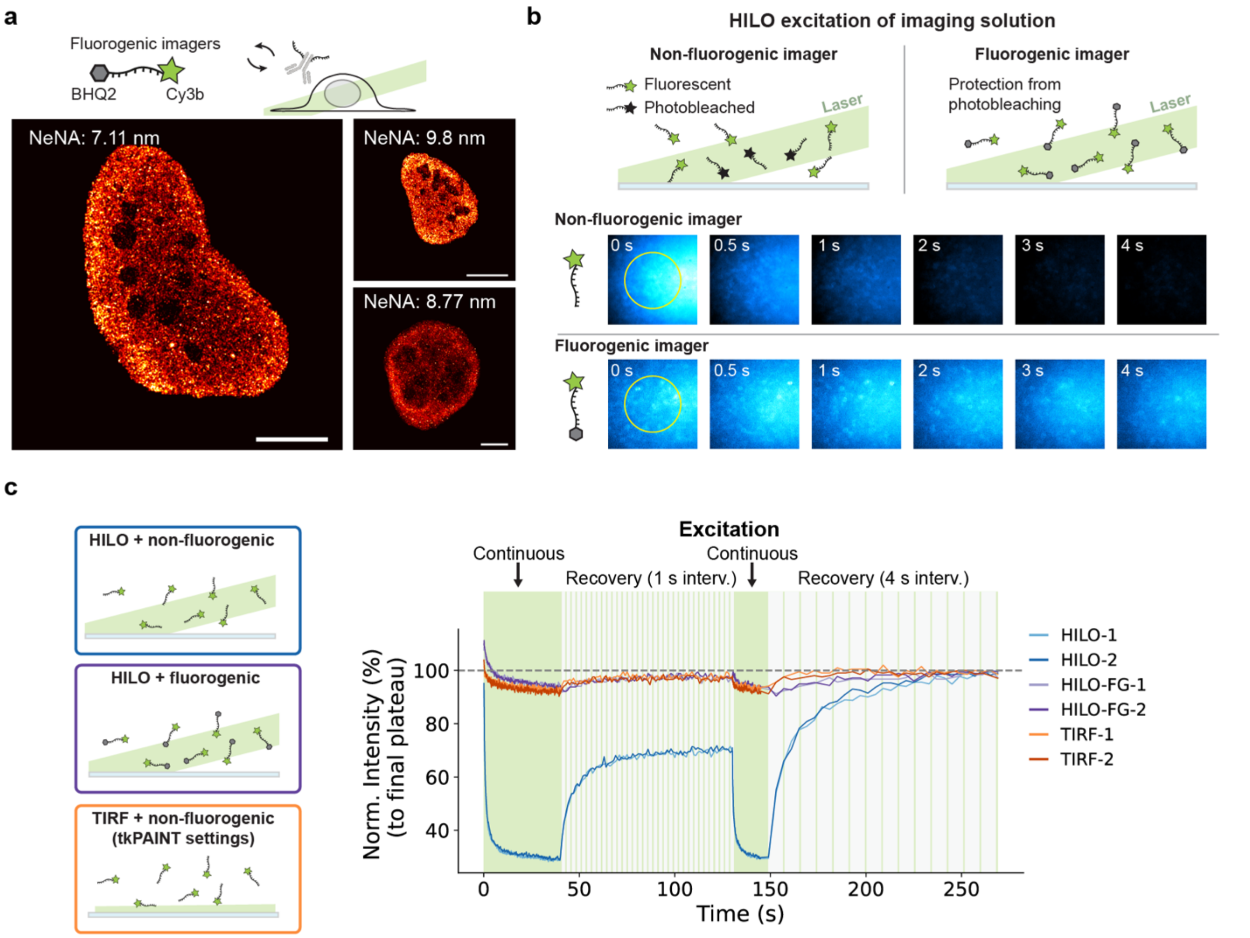


**Supplementary Fig. 3 | tkPAINT does not require fluorogenic imagers for bleaching protection. a**. HILO DNA-PAINT images of Pol II acquired using a fluorogenic imager design featuring a quencher molecule to suppress fluorescence of unbound imagers. The localization precision determined via NeNA is displayed for each dataset. **b**. Time series of imager bleaching upon HILO illumination. Fluorogenic imagers are protected from photobleaching in their unbound state during diffusion. **b** Fluorescence recovery after photobleaching experiment from the center circle as indicated by yellow circles in (b). First, the sample was continuously illuminated for 40 s and images acquired at 100 ms exposure time. Subsequently, fluorescence recovery was initiated by allowing 1 s intervals without excitation for 90 s. An additional 15 s period of bleaching was performed via full excitation before a second fluorescence recovery period was initiated allowing 4 s intervals without excitation for 120 s. Two independent repeats were performed for non-fluorogenic imager + HILO (blue), fluorogenic imager + HILO (purple) and non-fluorogenic imager + TIRF (the same acquisition settings as tkPAINT experiments; orange). Scale bars, 10 µm.

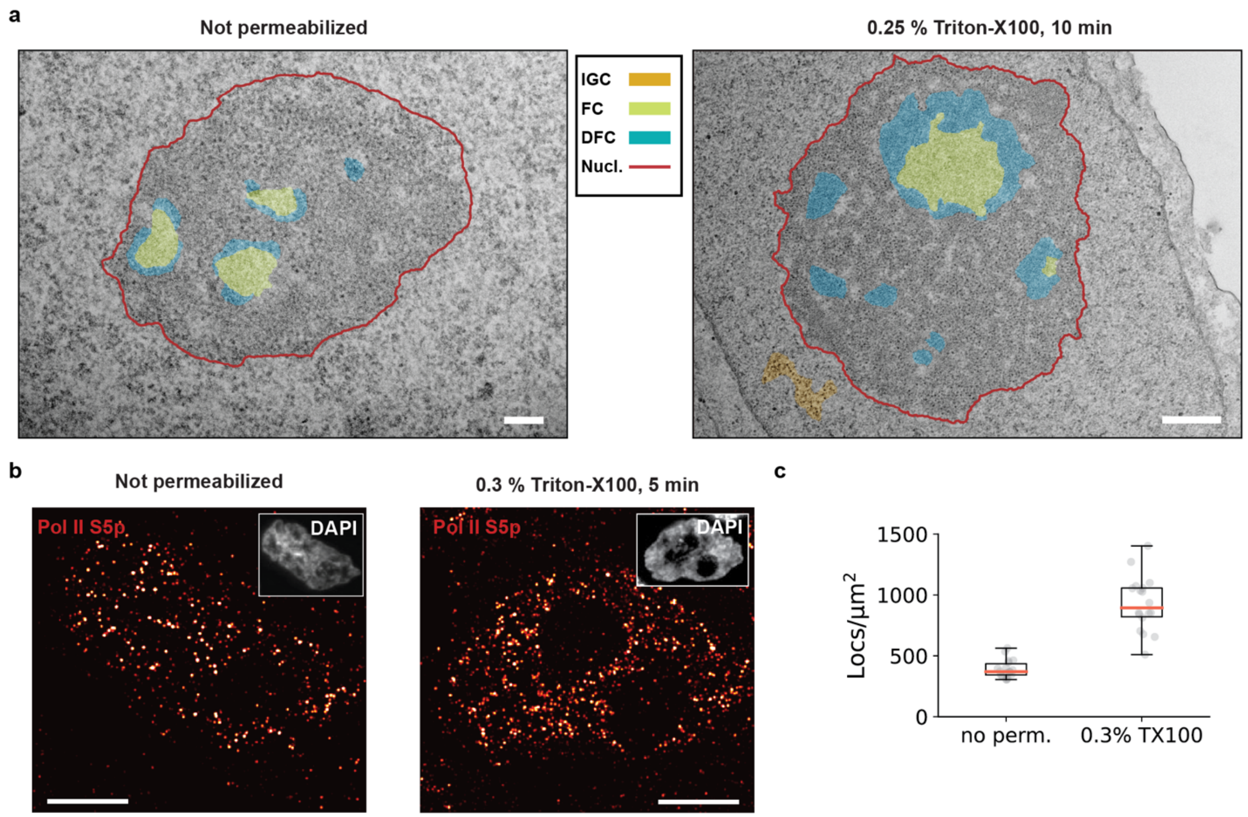


**Supplementary Fig. 4 | Permeabilization enhances nuclear epitope accessibility without altering nuclear ultrastructure.** **a** Annotated nuclear ultrastructure in TEM images shown in Extended Data Fig. 2a: 15 min fixation in 4 % PFA. No permeabilization (left image) vs. 10 min permeabilization in 0.25 % Triton X-100 (right image). Annotations: nucleolus (red line), dense fibrillar component (DFC, turquoise), fibrillar center (green) and interchromatin granular clusters (IGC; also nuclear speckles; yellow). Nuclear ultrastructure is relatively well-preserved even for such brief PFA fixation and permeabilization. **b** The Tokuyasu protocols developed by the Pombo lab (fixation: 4 % PFA for 10 min followed by 7 % PFA for 120 min) leverage brief permeabilizations with 0.2-0.3% Triton X-100 for 5-10 min[6,7] to enhance epitope access for immunolabeling throughout the section volume. In Supplementary Fig. 1 we validated via TEM that nuclear ultrastructure remains unaffected by this permeabilization step. Comparing tkPAINT images of Pol II S5p in non-permeabilized (left image) vs. permeabilized 150 nm-thin cryosections (0.3% Triton X-100 for 5 min; right image) visually confirmed increased labeling. The inset shows the DAPI channels of each nucleus. **c** Localizations per nuclear area measured for non-permeabilized vs. permeabilized tkPAINT datasets quantitatively confirming increased labeling. Due to higher antibody labeling, imager concentration in permeabilized tkPAINT datasets was half of that in non-permeabilized datasets and hence nuclear localization density was concentration adjusted. Scale bars, 200 nm and 600 nm in (a; left and right, respectively) and 3 µm in (b).

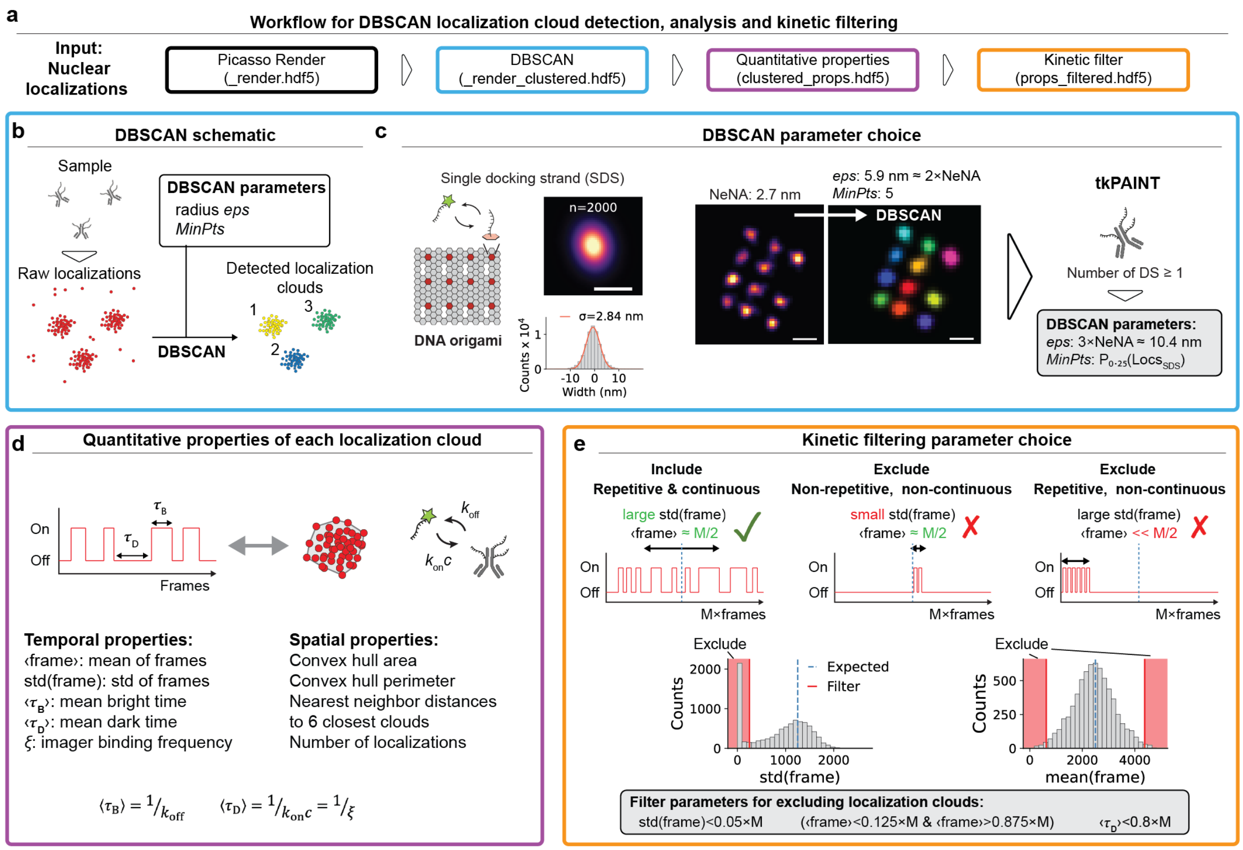


**Supplementary Fig. 5 | DBSCAN localization cloud detection, quantitative analysis and kinetic filtering. a** Overview of quantitative localization cloud analysis. Starting point of our spatiotemporal analysis were drift-corrected nuclear localizations (see **Supplementary Fig. 2**). Subsequently the clustering algorithm DBSCAN was applied to identify localization clouds (blue box). For each localization cloud, its spatiotemporal quantitative properties were calculated (purple box). Finally, a kinetic filter was applied to remove non-repetitive localization clouds (orange box). **b** DBSCAN schematic. DBSCAN has two input parameters[8], *eps* (or epsilon; distance between points to be considered as within neighborhood) and *minPts* (number of points required to form a dense region). Both parameters must be chosen such that neighboring localization clouds are registered as single clouds. **c** DBSCAN parameter choice. In theory, the NeNA localization precision should correspond to the standard deviation $\sigma$ of Gaussian distributed localizations from a single emitter[5] and has thus been suggested as input parameter for DBSCAN: *eps*=NeNA[9]. We first validated the relation between the global NeNA value of a dataset and $\sigma$ using DNA origami data acquired under tkPAINT conditions, where localizations from single emitters, i.e. single docking strands (SDSs) could be unambiguously identified. Indeed, NeNA and $\sigma$ were in close agreement. However, since $\sigma$ only contains ~68 % of Gaussian distributed data points, *eps* needs to be increased to capture also the outer localizations of a SDS localization cloud. On origami, we empirically found that adjusting *eps* to ~2xNeNA and *minPts* to 5 efficiently captured SDS localization clouds (colored SDSs on origami image indicate their identification as distinct clouds). In tkPAINT datasets, even a single primary antibody can yield overlapping localization clouds originating from more than one docking strand, because (i) an unknown number of docking strands is conjugated to each secondary antibodies or (ii) up to two nanobodies binding per primary antibody with one docking strand each. Since these localization clouds overlap, *eps* needs to be further increased. For tkPAINT we thus chose an average value corresponding to 3xNeNA as *eps* input. For *minPts* we chose the 25$^{th}$ percentile of the number localizations of localizations a SDS yielded on DNA origami imaged under identical conditions. **d** Calculation of quantitative spatiotemporal properties of each localization cloud. The left schematic shows a fluctuating intensity trace corresponding to a localization cloud with periods of imager binding ('on' or 'bright') and periods in between binding events without any localizations ('off' or 'dark'). The dwell times in each state are referred to as bright times $\tau_B$ and dark times $\tau_D$, respectively, and allow for calculating imager association and dissociation rates ($k_{on}$ and $k_{off}$, respectively) in DNA-PAINT[10] (see also **Extended Data Fig. 3** for a discussion on binding frequency $\xi$). Exploiting previously developed custom Python modules (picasso_addon[11], lbFCS[12] and lbFCS2[13]), we calculated the stated temporal properties (⟨frame⟩, std(frame), $\langle\tau_B\rangle$, $\langle\tau_D\rangle$) and the number of localizations for each localization cloud. Using Scipy[14], we further computed the convex hull area, perimeter as well as the nearest neighbor distances to the six closest localization clouds. **e** Kinetic filtering parameter choice. **Top:** three schematic fluctuating intensity traces for a tkPAINT data acquisition of *M* frames are shown. The left trace shows repetitive imager binding over the course of the experiment and thus has a mean frame ⟨frame⟩ value close to

$M/2$ (dashed line). Due to many binding events, the standard deviation of frames is relatively large. Such a trace would fulfill the kinetic filtering criteria. In contrast, the center and right intensity traces correspond to exemplary localization clouds not fulfilling the kinetic filtering criteria and are thus removed prior to further analysis. Bottom: Distributions of ⟨frame⟩ and std(frame) for a tkPAINT dataset of 5,000 frames length (left and right, respectively). The red shaded areas indicate localization clouds that were excluded through kinetic filtering. The blue dashed lines indicate the expected value for ⟨frame⟩ (i.e. at 5,000/2=2,500 for this dataset) and std(frame) (i.e. at ≈5,000/4=1,250 for this dataset). The following kinetic filtering cut-offs were used: (⟨frame⟩<0.125×M & ⟨frame⟩>0.875×M) and std(frame)<0.05×M. Furthermore, localization clouds with a mean dark time $\langle \tau_D \rangle$ of longer than 80 % of the acquisition length (i.e. $\langle \tau_D \rangle > 0.8 \times M$) were excluded.

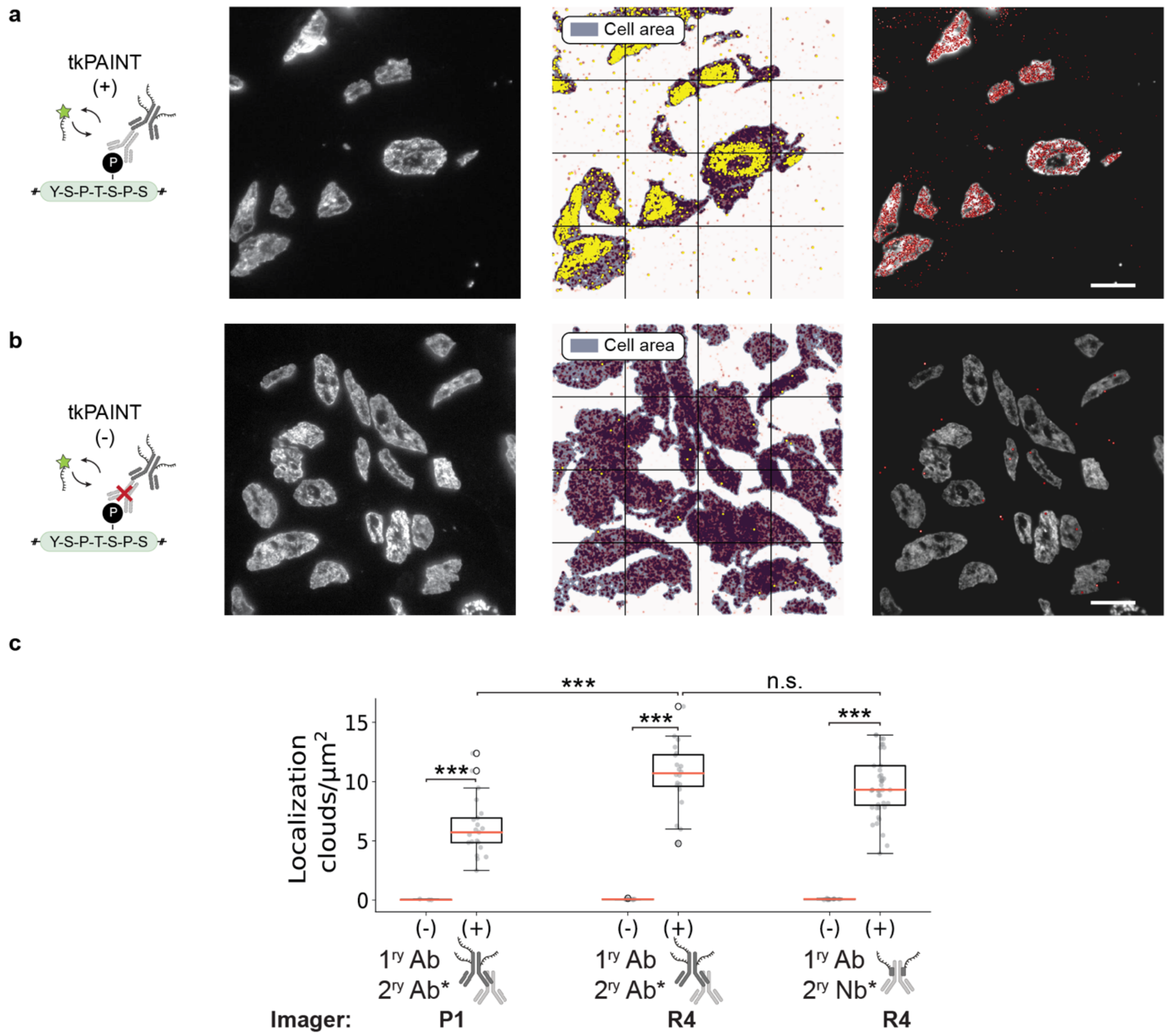


**Supplementary Fig. 6 | Kinetic filtering efficiently removes non-repetitive localization clouds and allows quantitative evaluation of labeling strategies. a** Exemplary tkPAINT Pol II S5p indirect immunolabeled (P1) dataset from **Fig. 2**a. Left: DAPI channel (left image). Middle: image containing all cellular localization clouds that passed kinetic filtering in yellow (both nuclear and cytoplasmic) overlayed over all recorded localizations marking the cellular area in blue. Right: All nuclear localization clouds that passed kinetic filtering (red). **b** Exemplary negative control tkPAINT dataset (P1) from **Fig. 2**c. Sample staining and imaging was performed as in a, but without addition of primary antibodies before incubation with DNA-conjugated secondary antibodies during sample preparation. Kinetic filtering efficiently excludes localization clouds that are not originating from primary antibodies. **c** Number of localization clouds per nuclear area for same imaging conditions as in **Fig. 2**b: (P1 – classic[1] vs. R4 – speed[15] using secondary antibodies) or comparing secondary labeling strategies (primary antibody (Ab) + secondary Ab (R4) vs. primary antibody + secondary nanobody (Nb)-R4). For each condition negative control (-) tkPAINT imaging was performed without the primary antibody but with the secondary label to assess the number of false positive localization clouds. False positives were negligible for all three cases (>140-fold difference between mean (+) and mean (-) for all three conditions). These tkPAINT datasets also allowed us to compare the detection efficiency between each labeling strategy, highlighting that secondary antibody-based tkPAINT imaging with speed imager R4 outperformed classic sequence P1, detecting 2× more nuclear Pol II S5p localization clouds (~12/µm2 vs. ~6/µm2, respectively). R4 in combination with secondary nanobodies yielded a similar density of nuclear localization clouds (~10/µm2), demonstrating that secondary antibody-based amplification is not required for efficient primary antibody detection in tkPAINT. Two-sample t-test: ***p ≤ 0.001, n.s. (not significant) p > 0.05. Scale bars, 10 µm.

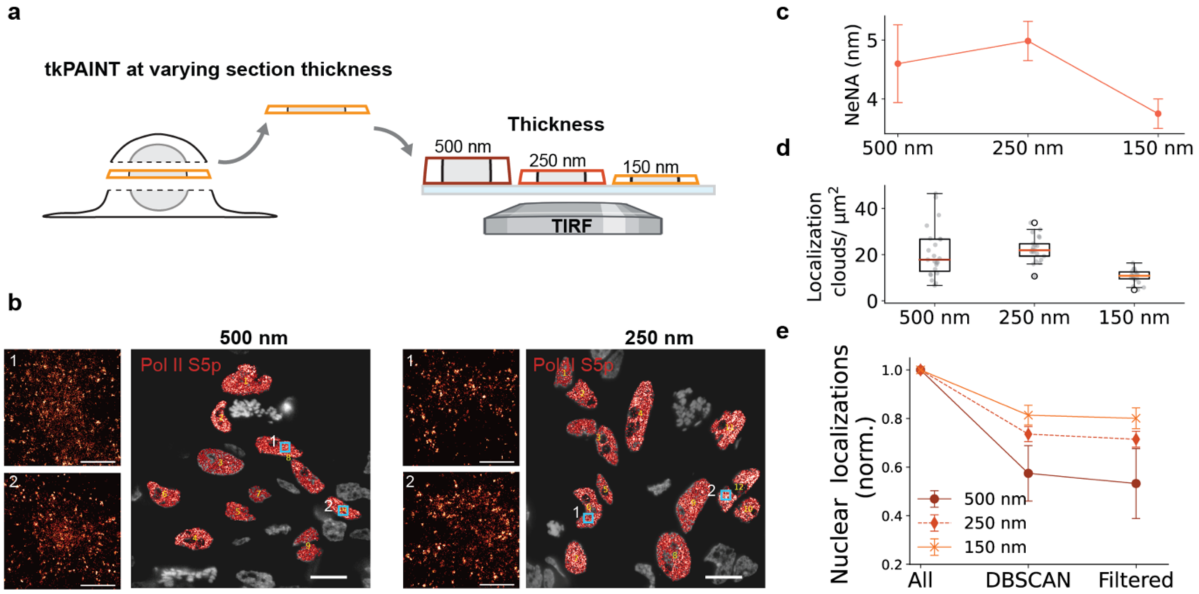


**Supplementary Fig. 7 | Influence of section thickness on tkPAINT imaging. a** Schematic illustrating tkPAINT imaging at three different cryosection thicknesses: 500 nm, 250 nm and 150 nm. **b** Exemplary tkPAINT S5p datasets shown for 500 nm and 250 nm cryosection thickness acquired under identical conditions as previous 150 nm thick cryosections (two datasets were acquired each). As expected from our discussion in Extended Data Fig. 3, the zoom-ins show that tkPAINT images become more crowded, since with increasing thickness more antibodies are present within the imaged volume. Note that we adjusted the TIRF angle such that we achieved roughly homogeneous illumination throughout 150 nm sections, i.e. reaching deeper into thicker sections, thereby increasing the imaged volume. This is most pronounced for 500 nm compared to 250 nm, as visible from many "false" localizations between localization clouds originating from simultaneously bound imagers[16] in the zoom-ins, which indicate that the imager concentration needs to be reduced. **c** NeNA localization precision decreases with cryosection thickness. Note that by adjusting the TIRF angle for minimal penetration depth, highest localization precisions should be achievable independent of section thickness[17]. **d** Number of localization clouds per nuclear area vs. cryosection thickness showing a linear increase from 150 nm to 250 nm, but then saturates for 500 nm thick cryosections. **e** Kinetic filter yield shown for tkPAINT Pol II S5p 500 nm, 250 nm and 150 nm datasets. Normalized localization counts with respect to all initial nuclear localizations showing relative loss of localizations in each analysis step. Scale bars, 10 µm.

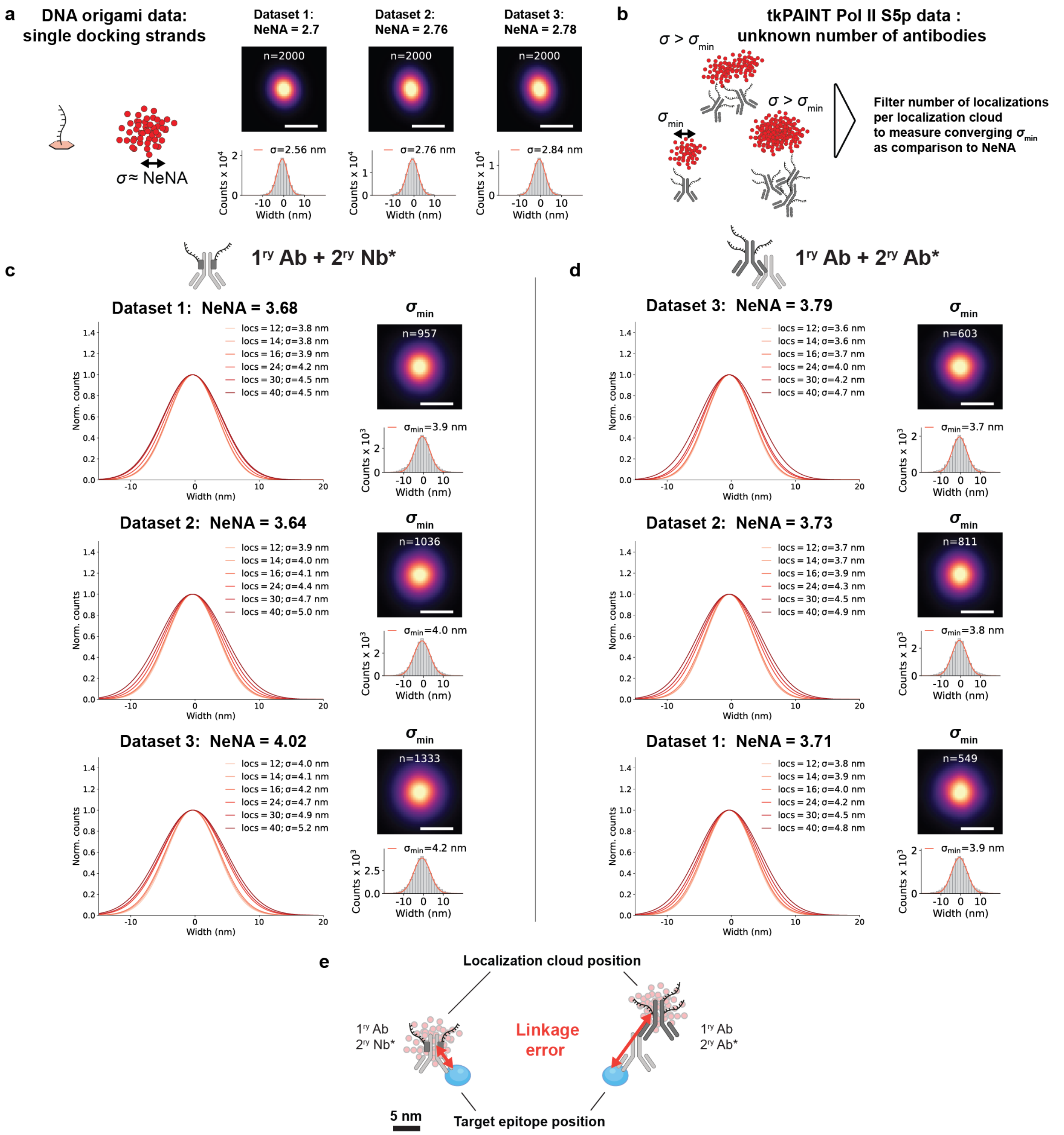


**Supplementary Fig. 8 | Scan for convergent minimum number of localizations per cloud – HeLa cells. a** Using DNA origami data acquired under identical conditions as tkPAINT experiments we measured the localization spread of single docking strands (SDSs). For each dataset we center-of-mass aligned 2,000 randomly-selected SDS localization clouds to obtain cloud average images (top). The histograms below show each localization distribution and a Gaussian fit (red curve) to obtain the standard deviation $\sigma$. The $\sigma$ of Gaussian distributed SDS localizations is in relatively close agreement with the global NeNA localization precision (2.8 ± 0.1 nm and 2.7 ± 0.1 nm, respectively; mean ± standard deviation. See also Supplementary Fig. 4). **b** In tkPAINT datasets, accumulations of antibodies with distances of only a few nanometers lead to enlarged localization clouds. We defined the standard deviation of the smallest localization clouds we could find as $\sigma_{min}$ ("minimum" standard deviation) and hypothesized these correspond to single antibodies. To test this, we performed a range of center-of-mass alignments for localization clouds filtering for different maximum number of localizations to find $\sigma_{min}$ as the converging standard deviation where further reduction in localization does not further reduce the localization spread. **c** Left: Range scan for maximum number of localizations per localization cloud for secondary nanobody-based tkPAINT Pol II S5p datasets to obtain the converging $\sigma_{min}$. Right: Localization distribution and Gaussian fit (red curve) for $\sigma_{min}$ as well as averaged sum images above. The number of localization clouds per sum image is stated above. **d** Same as c but for secondary antibody-based tkPAINT Pol II S5p datasets. **e** Schematic of linkage error. Although it was not possible to observe resolution differences in secondary-nanobody labeled primary antibodies as compared to DNA-conjugated secondary antibodies ($\sigma_{\mathrm{min}}$ of 4.0 ± 0.2 nm and 3.8 ± 0.1 nm, respectively), secondary nanobodies have the advantage of reducing the linkage error by which the physical size of the label displaces localization clouds away from the true epitope position. Observing similar $\sigma_{\mathrm{min}}$ can be explained by our range scan focusing on those secondary antibodies with only few docking strands, whereas for secondary nanobodies a maximum of two docking strands is present per primary antibody. Scale bars, 10 nm in averaged sum images.

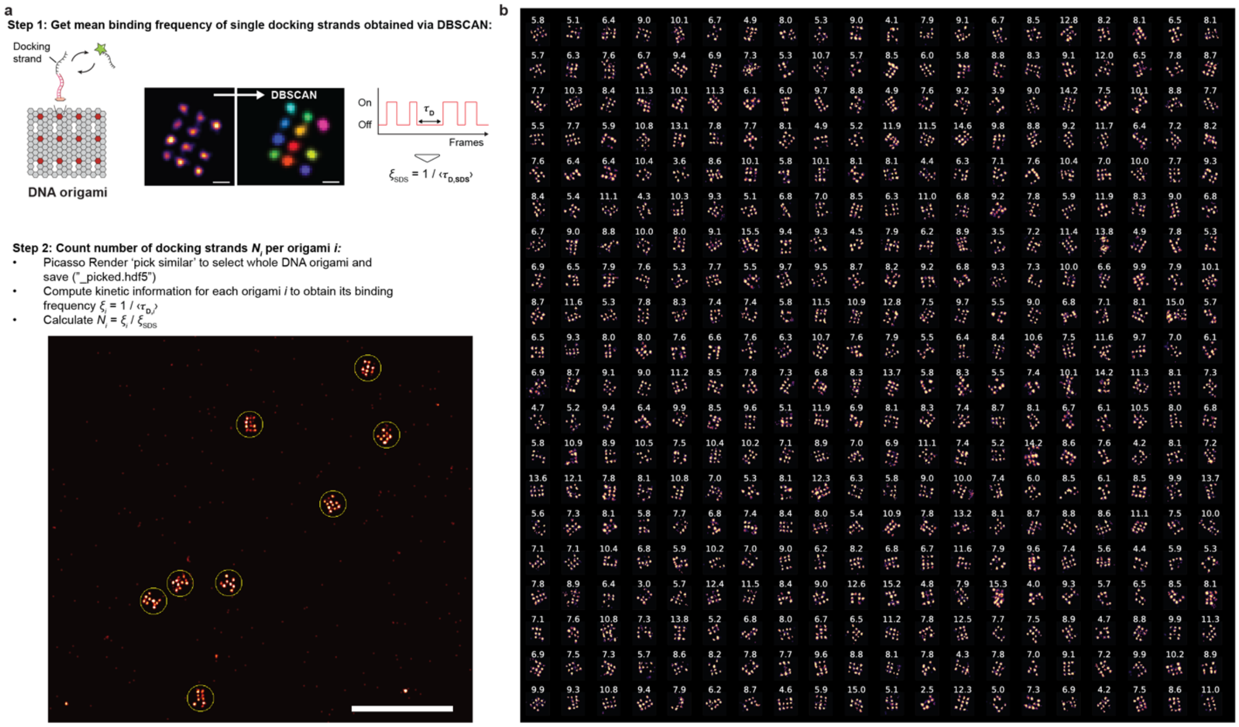


**Supplementary Fig. 9 | qPAINT analysis steps for DNA origami data. a** Analysis schematic for counting docking strands on DNA origami with qPAINT[1]. The origami design (same as in Extended Data Fig. 1) features binding sites to which up to 12 docking strand-adapter oligos can be stably hybridized. In step 1, we applied DBSCAN to identify single docking strands (SDSs) on DNA origami and computed the average SDS binding frequency $\xi_{\mathrm{SDS}} = 1/\langle \tau_{\mathrm{D,SDS}} \rangle$ through measurement of the average dark times over all SDSs in the dataset. In step 2, we used Picasso Render[1] and its tool 'Pick Similar' to select entire DNA origami (yellow circles) and computed the binding frequency of each individual DNA origami $\xi_i$. The qPAINT counting result for each origami is then obtained via $N_i = \xi_i / \xi_{\mathrm{SDS}}$. **b** 400 randomly selected DNA origami structures oriented as 20x20 grid. The qPAINT counting result is displayed above each origami. Scale bars, 500 nm in (b).

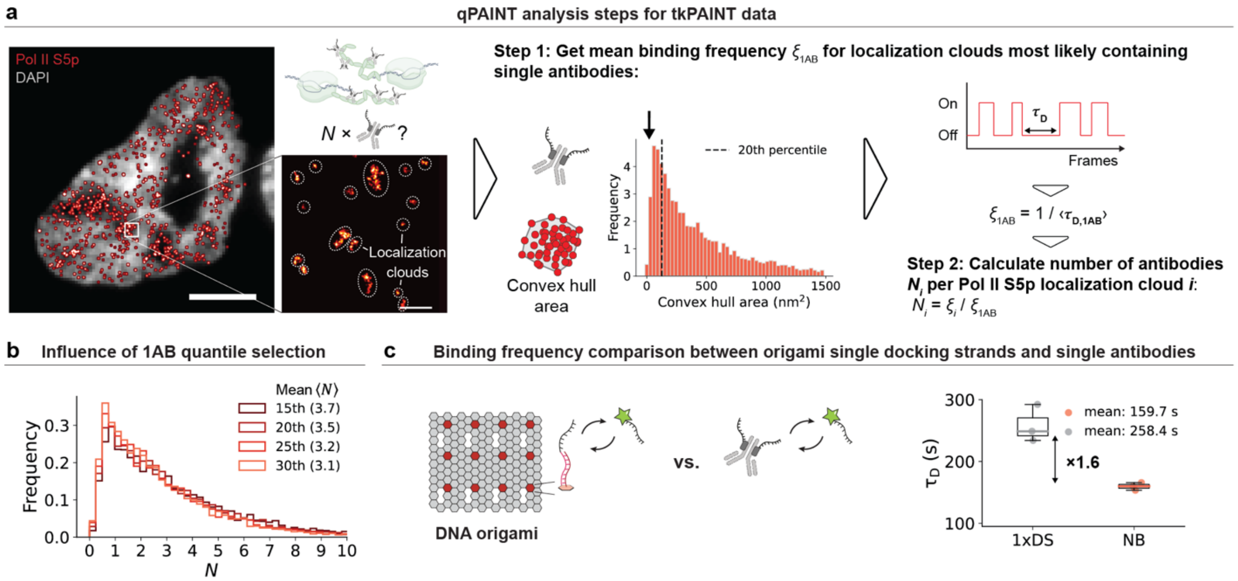


**Supplementary Fig. 10 | qPAINT analysis steps for tkPAINT data. a** Same Pol II S5p tkPAINT dataset as in Fig. 3a. Datasets featured sparse localization clouds from single antibodies and larger localization clouds containing several antibodies. In step 1 we defined a cut off at the 20th percentile of the convex hull area to define single antibody localization (1AB) clouds as qPAINT reference. For these, we calculated the reference binding frequency $\xi_{1AB} = 1/\langle \tau_{D,1AB} \rangle$ through measurement of the average dark times. In step 2, we obtained qPAINT antibody counting results for each localization cloud $i$ in the dataset via $N_i = \xi_i/\xi_{SDS}$. **b** The influence of the convex hull area cut off choice on final counting results was minor. **c** Comparison of the mean dark time between single docking strands on DNA origami and single antibody localization clouds revealed 1.6-fold reduction, indicating 1.6 docking strands per primary antibody (note that only up to two nanobodies can bind one primary antibody and each nanobody carries one docking strand). Scale bars, 3 µm in (**a**) and 1 µm in zoom-in.

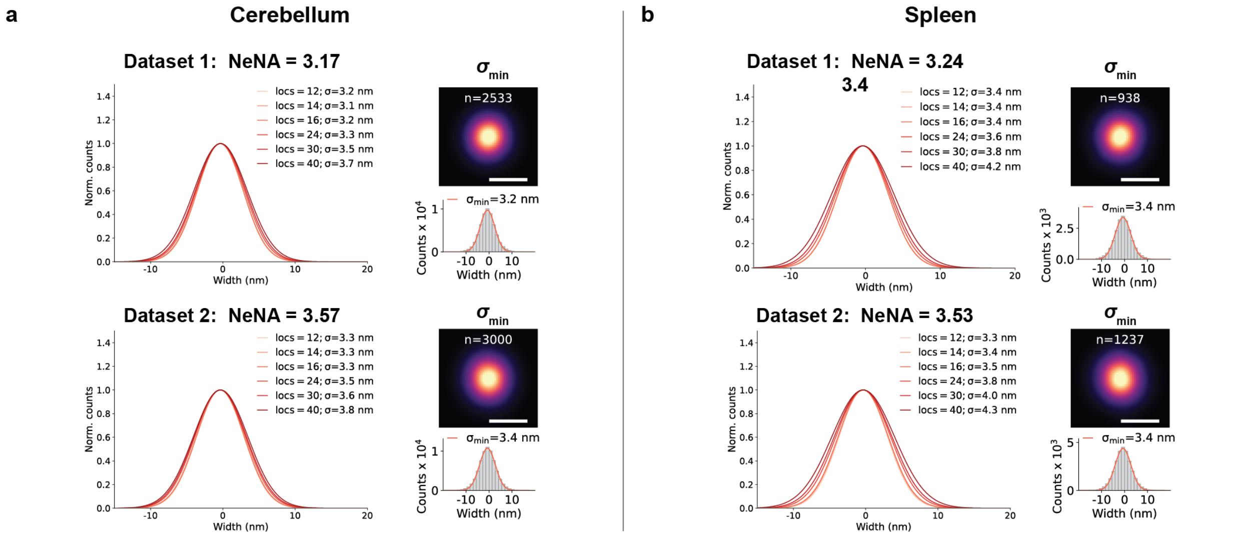


**Supplementary Fig. 11 | Scan for convergent minimum number of localizations per cloud – mouse tissues. a** This figure shows the same analysis introduced in Supplementary Fig. 8 for HeLa cells, but for the mouse cerebellum datasets shown in Fig. 4. Left: Range scan for maximum number of localizations per localization cloud for secondary nanobody-based tkPAINT Pol II S5p datasets to obtain the converging $\sigma_{min}$. Right: Localization distribution and Gaussian fit (red curve) for $\sigma_{min}$ as well as averaged sum images above. The number of localization clouds per sum image is stated above. **b** Same as a, but for mouse spleen datasets shown in Fig. 4. Scale bars, 5 nm.

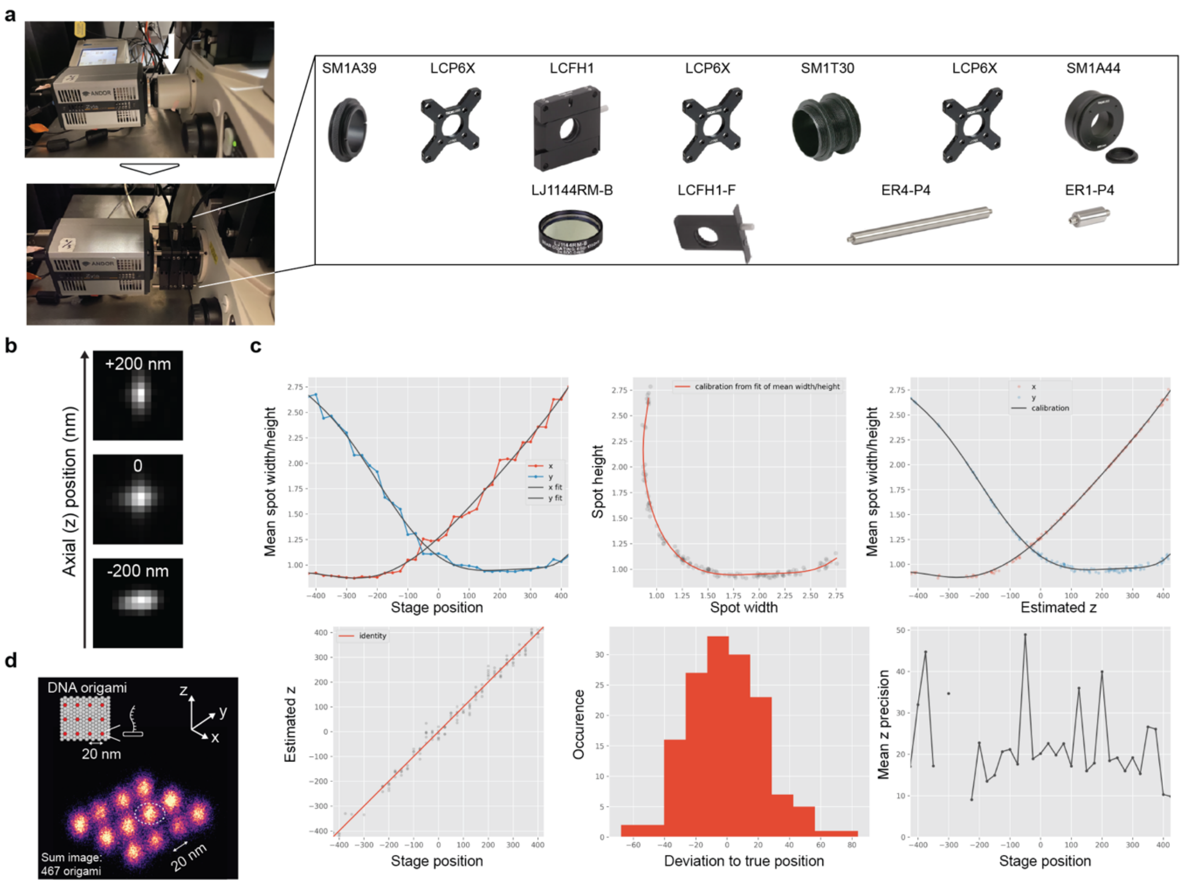


**Supplementary Fig. 12 | Custom cylindrical lens insertion for 3D tkPAINT. a** Parts list of Thorlabs optics components to replace the C-mount port (white arrow) at our standard Nikon TI Eclipse TIRF system with a cylindrical lens inset. Approximate cost at the time this manuscript was written: ~700 $. Part LCFH1-F can be purchased extra to leave an empty lens holder at the microscope for standard TIRF microscopy. The threaded tubing and our general design allow a flexible adjustment of the total distance between the camera and the port to match the focal length of the tube lens. **b** Axial calibration z-stack acquired using fluorescent beads. **c** Picasso Localize[1] output of 3D calibration: the z-stack was acquired at a step size of 25 nm and then loaded into Picasso Localize[1]. Running "Calibrate 3D", a box size large enough to fit the enlarged astigmatic point spread function was chosen. The generated 3D calibration .yaml file can be loaded when localization a 3D dataset, and the option "Fit z" needs to be ticket. Importantly, since insertion of an additional lens affects the magnification, this can be compensated for in Picasso. In our case, we determined the magnification factor by using DNA origami that carried a pre-designed 20 nm spacing as a nano ruler (**d**).

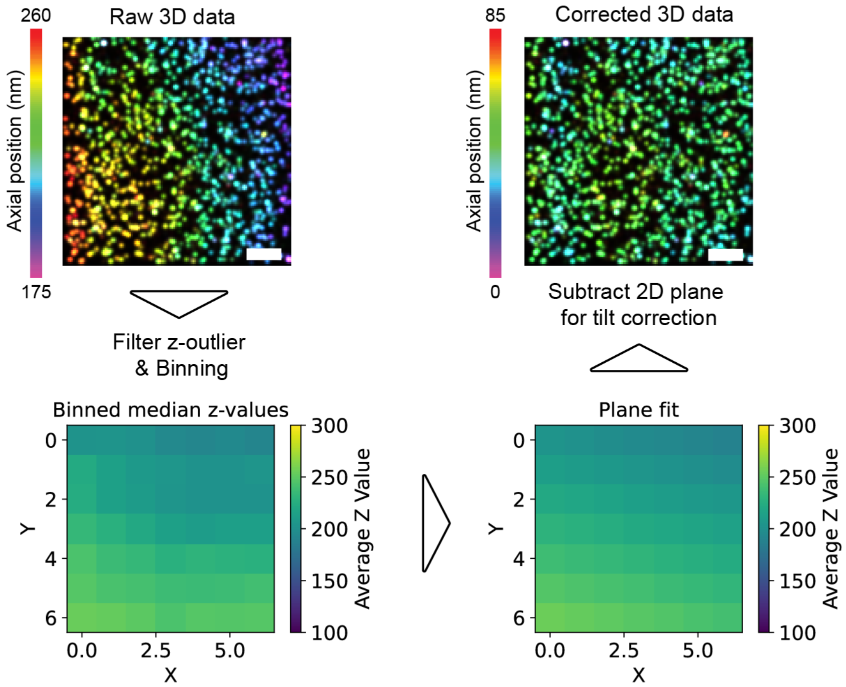


**Supplementary Fig. 13 | Planar fit for axial tilt-correction in 3D tkPAINT.** Axial tilt correction workflow shown at the example of a DNA origami 3D DNA-PAINT set using a custom Python implementation inspired by ref.[18]. After removal of z-outliers (>4x median) the 3D dataset was binned into a pixelated map where each pixel was assigned the median z-position of all localizations within the pixel. Next a 2D plane was fit to the pixel map to approximate any tilt. Lastly, the planar fit was subtracted from the initial 3D tkPAINT dataset to remove axial tilt and normalize to z=0. Scale bars, 5 µm.

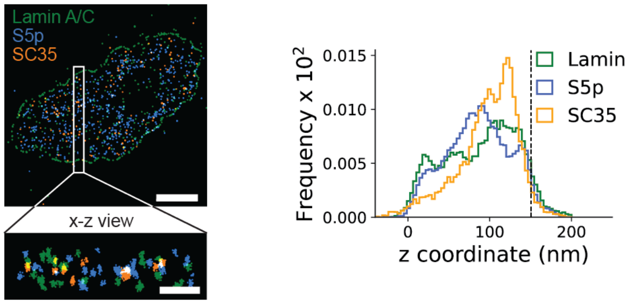


**Supplementary Fig. 14 | Axial distribution of antibody signal for 3D Exchange-tkPAINT.** Axial distribution of antibody signal for 3D Exchange-tkPAINT (Lamin A/C, POL II Pol II S5p and SC35) displayed in **Fig. 5**c. The dashed line indicates the set cutting thickness of 150 nm. Scale bars, 3 $\mu$m in (**a**) and 150 nm in zoom-in.

### Supplementary Table 1 | Imaging parameters for tkPAINT/DNA-PAINT

Available as MS Excel file.

### Supplementary Table 2 | Used DNA-PAINT sequences

| Imager-Docking ID | Docking sequence | Imager sequence |
|---|---|---|
| P1 | ttATACATCTA | CTAGATGTAT-Cy3b |
| Pm2 | ttTCTTCTTCTTCTTCTTCTTCTT CTTCTTCT | GAGGAGG-Cy3b |
| R2 | ttACCACCACCACCACCACCA | TGGTGGT-Cy3b |
| R3 | ttCTCTCTCTCTCTCTCTCTC | GAGAGAG-Cy3b |
| R4 | ttACACACACACACACACACA | TGTGTGT-Cy3b |
